# White Matter Myelin Shapes Macroscale Functional Connectivity Through Integrative Communication

**DOI:** 10.64898/2026.03.22.713515

**Authors:** Mark C. Nelson, Wen Da Lu, Ilana R. Leppert, Golia Shafiei, Heather A. Hansen, Christopher D. Rowley, Bratislav Misic, Christine L. Tardif

## Abstract

White matter structural connectivity constrains large-scale brain communication, yet most network models do not account for biologically meaningful differences between connections. Although axonal diameter and myelination influence neural signaling at the microscale, how these features shape systems-level functional connectivity remains unclear. Here, we test whether structural connectomes weighted by white matter microstructure give rise to distinct communication regimes that differentially predict multimodal functional connectivity. Combining quantitative MRI and advanced diffusion modeling, we constructed whole-brain networks weighted by tract caliber and multiple myelin-sensitive measures. To these, we applied routing- and diffusion-based communication models and used the resulting communication metrics to predict haemodynamic and frequency-resolved electromagnetic connectivity. Myelin-weighted networks preferentially enhanced long-range communication efficiency and redistributed spectral energy toward globally integrative topological eigenmodes. In contrast, caliber-weighted networks emphasized mesoscale organization and short-range communication. Across nested regression models controlling for geometric embedding and network topology, myelin-sensitive communication explained unique variance in functional connectivity with effects varying systematically across cortical systems and frequency bands. The strongest coupling was observed for alpha-band connectivity in association and attentional networks, consistent with a role for myelin-dependent communication delays in supporting long-range alpha synchrony. These findings demonstrate how distinct white matter microstructural features give rise to heterogeneous large-scale communication regimes: tract caliber and myelin bias communication toward locally specialized and globally integrative architectures, respectively. By integrating biologically informed connectomics with communication modeling and multimodal functional data, this work advances a mechanistic account of how white matter microstructure shapes macroscale brain dynamics.

## Introduction

White matter connectivity constrains how signals propagate between brain regions and provides the anatomical substrate for large-scale functional interactions^1^. Yet structural connectivity does not uniquely determine functional connectivity (not a 1-1 mapping)^2^, and strong functional coupling can arise between regions that are only weakly or indirectly connected^3,4^. Most studies linking structure and function rely on streamline count or binary representations of white matter connectivity(Liu et al., 2023; Suárez et al., 2020), implicitly treating white matter as a uniform medium. As a result, the biological features of white matter that shape large-scale communication and functional coupling remain incompletely understood.

White matter is biologically heterogeneous, with axons varying in properties that are directly relevant for neural signaling^6^. At the microscale, the cross-sectional diameter of an axon contributes to the capacity, reliability, and conduction velocity of signal transmission^7^. The extent of axonal myelination also modulates conduction velocity while supporting a range of functionally relevant roles, including temporal precision, trophic support, metabolic efficiency, and activity-dependent plasticity^8,9^. Although these properties vary systematically across the brain and operate at different spatial and temporal scales^10,11^, their systems-level roles are not well established as they are rarely incorporated into whole-brain network models.

Increasingly, studies have begun to address this gap by integrating biologically informed white matter properties into structural connectomes^12,13^. Adopting this biologically-enriched perspective reframes white matter connectivity as a dynamic contributor to both network organization^14,15^ and cognition-related plasticity^16^. While we previously used this approach to demonstrate enhanced structure-function coupling^17^, these relationships were assessed via direct edgewise mappings and therefore could not incorporate multi-step network interactions. Given that most functional connections arise between indirectly connected regions^3^, modeling signal propagation across intermediate pathways becomes essential for understanding structure–function coupling at the systems level.

Network communication models^18^ offer a useful framework for bridging structural connectivity and functional interactions. These models formalize how signals may traverse a network, providing mechanistic abstractions of interregional communication rather than literal descriptions of neural processes. Importantly, the dynamics captured by these models are not tied to a single temporal scale and may therefore relate differently to neural interactions measured across distinct functional modalities.

Broadly, communication models can be arranged along a spectrum^19^: from routing-based models (e.g., ^4^), which emphasize selective and path-dependent signal transmission; to diffusion-based models (e.g., ^20^), which capture distributed and redundant signal spreading. Previous work has shown that different communication models can help explain both functional connectivity and behavior^4,21^. However, most studies fix the underlying structural weights and vary the communication rule, leaving open how the biological attributes of white matter interact with communication dynamics^22^.

In this study, we integrate biologically informed white matter connectivity with network communication modeling to examine how white matter features shape structure–function coupling. We estimate structural connectivity networks weighted by myelin-sensitive and caliber-sensitive measures at the tract-level. These are used to quantify both routing-based and diffusion-based communication patterns, which are then related to both haemodynamic and frequency-resolved electromagnetic connectivity. Rather than focusing on a single optimal model^23^, we ask how explanatory power is redistributed across communication regimes, topological scale, and cortical hierarchy. We find that myelin-weighted networks preferentially support globally integrative communication, characterized by long-range interactions and strongest coupling to alpha-band functional connectivity. In contrast, tract caliber-weighted networks emphasize spatially local, mesoscale organization. Together, these results support a view in which heterogeneous white matter structure gives rise to distinct neural signaling protocols, shaping the frequency-specific organization of functional connectivity across brain regions.

## Results

Analyses were performed using a multi-modal MRI dataset acquired at 3 tesla in 30 healthy adults (14 men, 16 women; 29±6 years of age). Structural connectivity networks were derived from diffusion and magnetization transfer (MT) MRI. The in-sample functional connectivity (FC) network was derived from resting-state blood oxygen-level-dependent functional MRI (BOLD-fMRI) as the cross-correlation of node-wise time series (**BOLD_in_**). An additional seven out-of-sample FC networks were obtained in fully preprocessed^24^ and group-averaged format. One was derived from resting-state fMRI (**BOLD_out_**), and six were derived from resting-state magnetoencephalography (MEG) corresponding to the canonical electrophysiological frequency bands: **delta** (2-4Hz), **theta** (5-7Hz), **alpha** (8-12Hz), **beta** (15-29Hz), **gamma_lo_** (30-59Hz), and **gamma_hi_** (60-90Hz). These data were collected in healthy participants (n = 33; age range 22 to 35 years) and made openly available through the Human Connectome Project (HCP; S900 release(Van Essen et al., 2013)). All networks were constructed at the whole-brain, cortex level using the Schaefer-400 atlas^25^ and served as the substrate for subsequent topological, communication, and structure–function coupling analyses.

We examined several weighted structural connectivity matrices that differed in the edge attribute used to parameterize interregional connections. Using advanced diffusion modeling^26,27^, a tract caliber–weighted network was estimated, which quantifies the *total cross-sectional area of the intra-axonal compartment* for each bundle of streamlines representing a network edge. Note that this network quantifies caliber at the edge- or tract-level and should not be confused with measures of axon-caliber^28^. We also considered multiple myelin-sensitive weightings, including: MTsat as a measure of myelin density, g-ratio as a measure of myelin sheath thickness relative to axon diameter, and tract delay as a measure of myelin-dependent conduction delays. The delay for each tract was estimated using its g-ratio, length, and a simplified estimation of conduction velocity. These edge weightings—caliber, myelin density, g-ratio, and delay—provide complementary perspectives^13^ of white matter microstructure and geometry while preserving a common network topology.

Throughout the Results, tract caliber is treated as a reference structural network, reflecting its conceptual similarity to widely used streamline-based connectomes^13^. Differences observed under myelin-sensitive weightings are therefore interpreted as reflecting the influence of microstructural and temporal constraints beyond those captured by tract caliber alone.

These analyses require both weight (high values reflect strength) and cost (low values are more optimal) representations for all structural networks. Where necessary in topology and communication analyses, tract delay was converted to a strength representation (1/delay), such that larger weights reflected stronger or more efficient communication. These will be differentiated as *delay* and *1/delay* throughout.

The Results are organized as follows: (i) descriptive characterization of myelin-weighted network trends, (ii) mesoscale topology, (iii) communication model behavior, and (iv) global and regional coupling with functional connectivity across timescales.

### Myelin-sensitive networks show divergent edge-weight alignment and convergent clustering

To contextualize subsequent modeling results, we first examined group-level relationships among white matter edge microstructural features, geometric properties, and BOLD functional connectivity, as well as their large-scale topological organization (Fig. 1).

**Figure 1.**
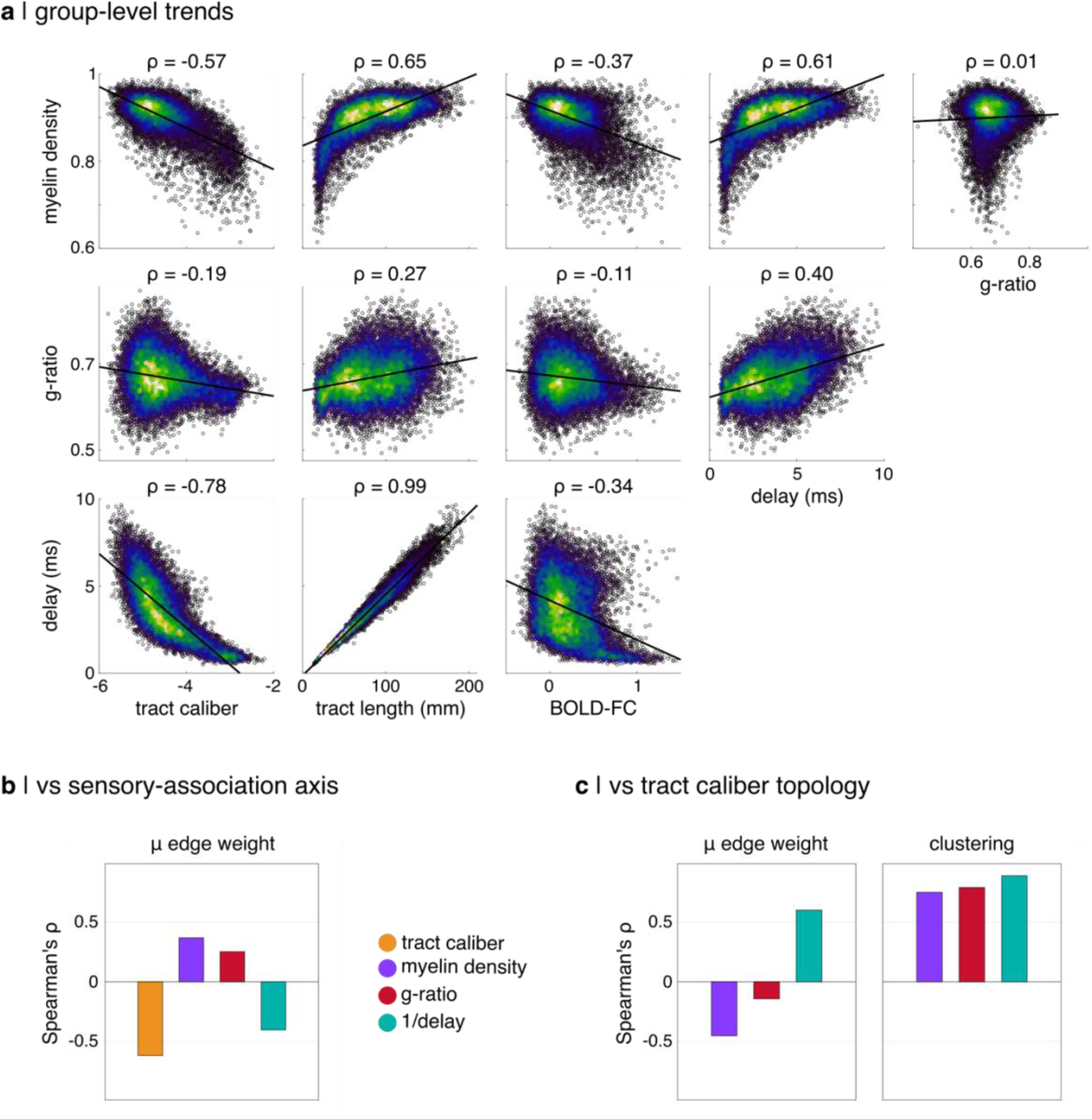
Group-level data trends and high-level topology. (**a**) Scatter plots show edge weights and Spearman’s rank correlations (ρ). The line represents best linear fit. Data is colored by density. (**b**) Nodewise Spearman’s rank correlations of mean edge weight with the sensory-association (S-A) axis ordinal mapping. (**c**) Nodewise Spearman’s rank correlation of mean edge weight and clustering coefficient for all myelin-weighted networks vs tract caliber. Tract caliber was derived using advanced diffusion modeling and myelin density from tractometry using MTsat images. Delay was derived from tract measures of g-ratio, length, and a simplified estimation of conduction velocity. For topology measures, delay was converted to 1/delay to align with other structural network weights.

#### Group-level edgewise data trends

Across all edges, myelin-related metrics exhibited systematic relationships with tract caliber, tract length, and BOLD functional connectivity (Fig. 1A). All myelin metrics were negatively correlated with caliber, with the strongest association observed for delay and the weakest for g-ratio. Conversely, all myelin metrics were positively correlated with tract length, again with delay showing the strongest dependence and g-ratio the weakest. All myelin metrics were negatively correlated with BOLD-derived functional connectivity. Myelin density (MTsat) and delay exhibited comparable effect sizes, and g-ratio again showed the weakest association. Delay was more strongly correlated with myelin density than with g-ratio, consistent with the strong influence of tract length on delay estimates in our implementation. Surprisingly, myelin density and g-ratio were uncorrelated at the edge level.

Together, these relationships indicate that, while the myelin-sensitive metrics share broad trends with geometric and functional properties, myelin density and g-ratio capture partially distinct aspects of white matter microstructure. This supports their joint use in predicting functional connectivity without exacerbating collinearity in multivariate models.

#### Nodewise gradients and spatial organization

At the node level, mean edge weight varied systematically along the sensory–association (S–A) axis^29^ (Fig. 1B). Mean myelin density and g-ratio were positively correlated with the S–A axis (ρ = 0.37 and ρ = 0.25, respectively), whereas 1/delay and caliber showed negative correlations (ρ = −0.40 and ρ = −0.62, respectively). This pattern indicates that association cortex is preferentially characterized by edges with higher myelin density and g-ratio, but lower caliber and slower transmission properties. This divergence along the S–A axis suggests that microstructural and geometric features encode complementary aspects of cortical hierarchy, rather than reflecting a single shared gradient.

#### Topological alignment with caliber

We next examined how myelin-related metrics align with tract caliber-derived network topology (Fig. 1C). The nodewise correlation of mean edge weight with caliber was strongly negative for myelin density, weakly negative for g-ratio, and strongly positive for 1/delay. In contrast, node clustering coefficients were strongly positively correlated with caliber across all myelin metrics. These results reveal a dissociation between local edge strength and higher-order network organization: while myelin-sensitive edge weights diverge from caliber, their induced topological structure substantially overlaps, particularly with respect to clustering.

Collectively, these results provide evidence for (i) broad similarity across myelin-sensitive metrics, (ii) partial divergence between myelin density and g-ratio at the level of raw data trends alongside convergence in network topology, and (iii) a relatively close alignment between tract caliber and 1/delay, particularly in their large-scale organizational signatures. These observations motivate treating myelin metrics as related but non-redundant constraints on network communication, rather than interchangeable proxies.

### Community structure

To examine how microstructural weighting shapes large-scale network organization, we next compared the community structure of tract caliber, myelin-sensitive, and timing-based structural networks (Fig. 2).

**Figure 2.**
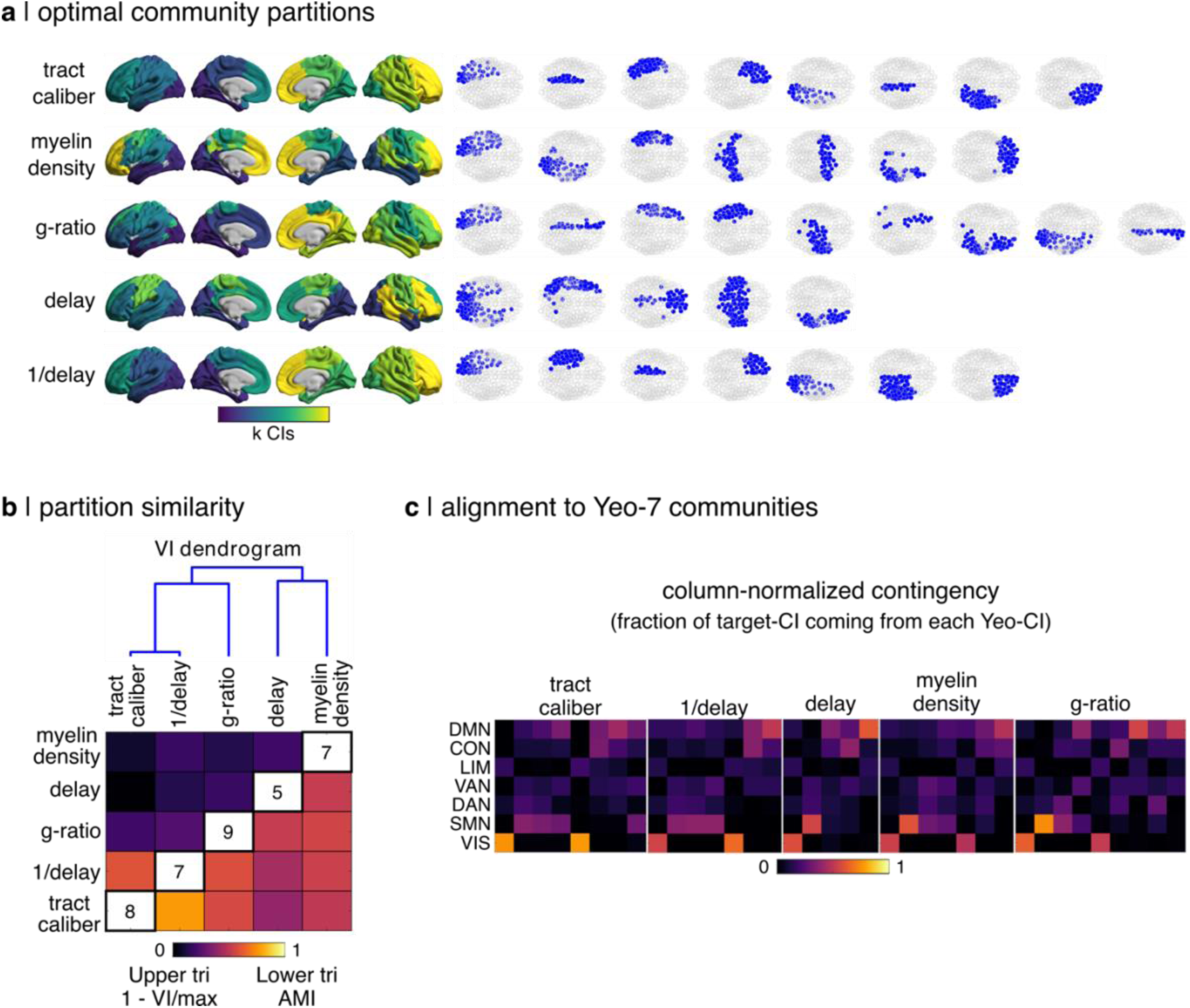
Community structure. (**a**) Optimal community partitions are shown on the cortical surface and in an axial view for all structural networks. Surface plot colors reflect integer values (k) for each community. (**b**) Partition similarity was quantified using adjusted mutual information (AMI) and variation of information (VI). The heatmap shows AMI on the lower triangle, 1 – VI / VI_max_ on the upper triangle, and k community counts on the main diagonal. The dendrogram was constructed using VI as a distance metric. (**c**) Heatmaps show the column-normalized contingency of each community with the 7-network Yeo atlas. Communities have been aligned across plots using a Hungarian matching algorithm to aid visualization. The Yeo networks correspond to: visual (VIS), somatomotor (SMN), dorsal attention (DAN), salience ventral attention (VAN), limbic (LIM), fronto-parietal control (CON), default mode (DMN).

#### Optimal community partitions

Optimal community partitions revealed marked differences in the spatial localization and symmetry of network modules across structural weightings (Fig. 2A). Tract caliber exhibited spatially localized communities (k = 8) that were perfectly mirrored across the midline, yielding four bilateral pairs corresponding to fronto-central, temporo-parietal, medial parietal, and occipital systems. Myelin density-based modules (k = 7) resembled tract caliber in posterior cortex but diverged in central and frontal regions, where communities were less spatially localized and frequently spanned both hemispheres. Notably, myelin density recovered separate modules resembling primary motor (M1) and primary somatosensory (S1) cortex. In contrast, g-ratio modules (k = 9) showed reduced posterior–anterior localization relative to tract caliber but did not exhibit midline-spanning communities, instead maintaining hemispheric separation. Delay-based modules (k = 5) displayed the least spatially localized community structure, characterized by a prominent sensorimotor-like community encompassing both M1 and S1, as well as posterior visual-like and frontal communities that spanned the midline. Modules for 1/delay (k = 7) closely resembled tract caliber, with the primary distinction being the lack of separation between medial and lateral parietal communities in the right hemisphere.

Overall, these patterns indicate a graded shift from spatially compact, hemispherically-segregated communities toward increasingly diffuse and midline-spanning organization across tract caliber, myelin-sensitive, and timing-based networks.

#### Partition similarity across structural networks

To quantify similarities among community partitions, we computed variation of information (VI) and adjusted mutual information (AMI) between all pairs of partitions (Fig. 2B). The resulting dendrogram revealed a relatively close alignment between tract caliber and 1/delay networks, as well as between delay and myelin density networks. g-ratio occupied an intermediate position, clustering more closely with tract caliber and 1/delay than with delay and myelin density. This organization mirrors the edge-level correlations observed in Figure 1 and suggests that similarities in raw microstructural trends propagate upward to mesoscale network organization.

#### Alignment with canonical functional systems

We next assessed the correspondence between structural communities and canonical functional systems using column-normalized contingency with the Yeo-7^30^ resting-state networks (Fig. 2C). The strongest correspondence with the visual network was observed for tract caliber, although this community was split across hemispheres. Sensorimotor systems were most strongly recovered in g-ratio-weighted networks, followed by myelin density and delay. The lower contingency observed for myelin density in this system likely reflects the splitting of M1 and S1 into separate communities. Among higher-order systems, the default mode network showed the strongest correspondence with delay-based communities. In contrast, limbic, ventral attention, and dorsal attention networks were not strongly represented in any of the structural weightings examined. These results indicate that structural community architecture preferentially aligns with unimodal functional systems, while higher-order association networks are weakly captured at the level of modular organization of static structural networks.

These results demonstrate both overlap and divergence in mesoscale organization across structural weightings. This variation in community organization provides a plausible substrate for strategy-dependent sensitivity of communication models to structural weighting, with mesoscale modular structure expected to differentially shape distinct communication processes.

### Communication model behavior across structural weightings

We next examined how the microstructural properties of network edges shape inter-regional communication patterns. We focus on pairwise comparisons with tract caliber as a reference, and we include six communication models covering the routing-diffusion spectrum: shortest paths efficiency (SPE), navigation efficiency (NE), search information efficiency (SIE), path transitivity (PT), communicability (CMY), and diffusion efficiency (DE) (Fig. 3). Routing-based models (SPE, NE) capture communication along specific paths through the network, whereas diffusion-based models (CMY, DE) reflect distributed signal propagation that depends on broader network organization^18,31^. Comparing these complementary strategies allows us to assess whether microstructural weighting differentially constrains local versus distributed modes of communication.

**Figure 3.**
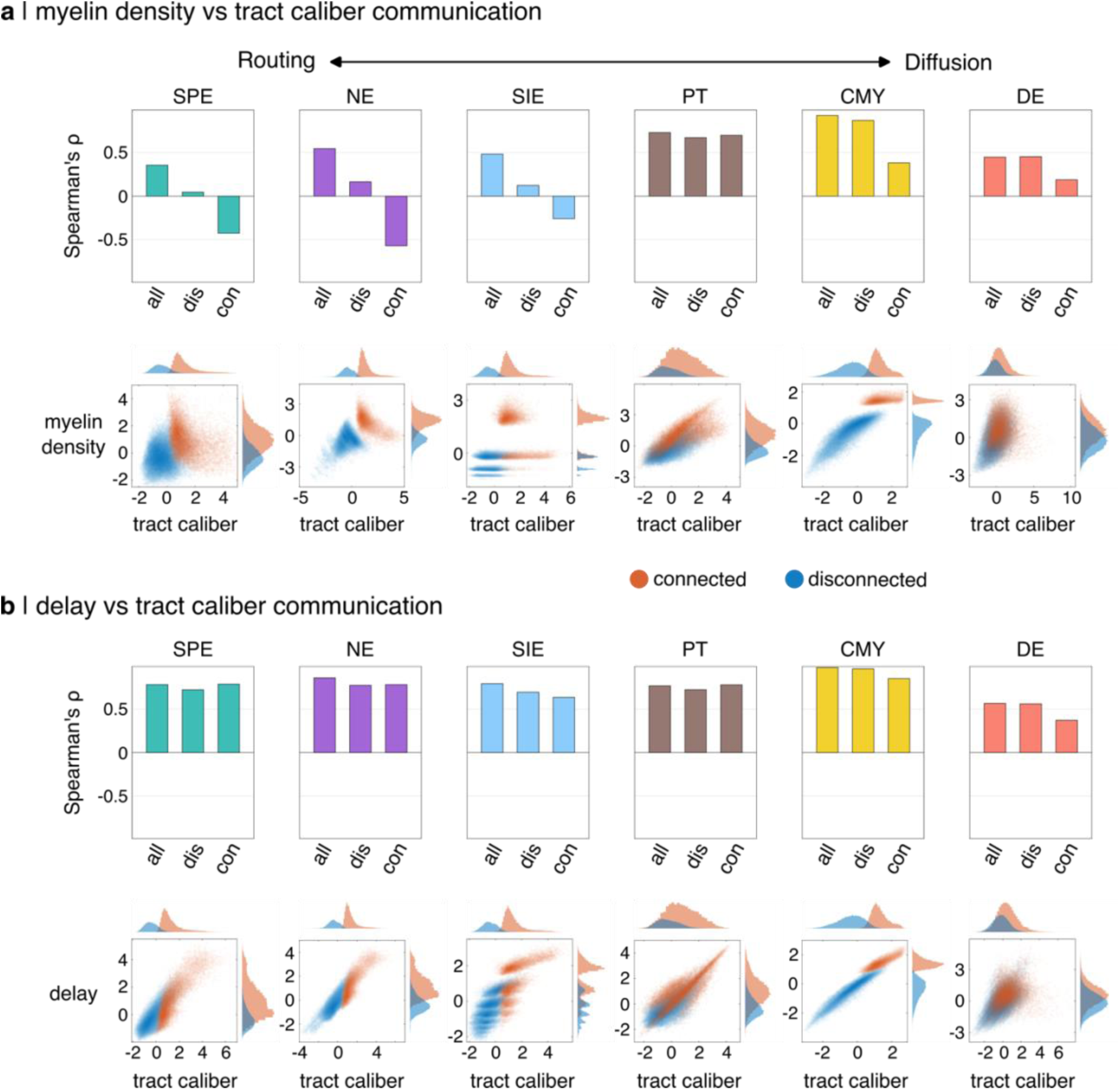
Communication model behavior across structural weightings. Six communication models were computed from all structural networks: shortest path efficiency (SPE), navigation efficiency (NE), search information efficiency (SIE), path transitivity (PT), communicability (CMY), and diffusion efficiency (DE). Edgewise communication values are compared with tract caliber for (**a**) myelin density and (**b**) delay. Data points in scatter plots are colored to reflect the presence (red) or absence (blue) of a direct structural connection, and color intensity reflects data density. Axis-aligned histograms show the distributions of both data groups. Bar plots represent the edgewise Spearman’s rank correlations of communication measures for three data groupings: all edges, disconnected edges (blue data points in scatter plot), and connected edges (red data points).

Communication models are designed to operate on specific representations of the input structural connectome: cost for SPE, NE, and PT; weight for CMY and DE; and both for SIE. Thus, the distinction between delay and 1/delay is not required from here.

Results for g-ratio–weighted networks are not shown, as they closely mirrored those obtained for myelin density across communication models.

#### Myelin density versus tract caliber communication

For routing-based communication, networks weighted by tract caliber and myelin density exhibited a strong inverse relationship for node pairs connected by a direct structural edge (Fig. 3A). In contrast, for node pairs without a direct structural connection, routing-based communication values were weakly correlated or uncorrelated across the two weightings. Diffusion-based communication showed a complementary pattern. For node pairs without a direct structural connection, diffusion measures derived from tract caliber and myelin density were strongly positively correlated, whereas correlations were weaker for node pairs with a direct connection. This dissociation indicates that routing-based communication is most sensitive to differences in edge weights along directly connected pathways, while diffusion-based communication reflects broader network organization that extends beyond direct structural links.

#### Delay versus tract caliber communication

In contrast to myelin density, delay-based communication closely aligned with tract caliber across all communication models (Fig. 3B). Strong positive correlations were observed between delay- and tract caliber-weighted communication values for both directly connected and indirectly connected node pairs, independent of communication strategy. This convergence suggests that delay and tract caliber impose similar constraints on both local and distributed communication processes, consistent with their close alignment in edge-level trends and mesoscale topology.

These results demonstrate that the impact of structural weighting on network communication depends jointly on the communication strategy and on whether node pairs are directly connected. Myelin density-weighted networks diverge most strongly from tract caliber under routing-based communication, particularly for directly connected edges, whereas delay-weighted networks show consistent alignment with tract caliber across strategies and connection types. Without implying that any single communication strategy is uniformly more sensitive to mesoscale organization, these results suggest a graded transition in the relative influence of edge weights versus global network structure on communication, which is aligned with the routing–diffusion spectrum.

### Functionally relevant differences between tract caliber and myelin communication

To assess whether differences in structural weighting translate into functionally meaningful distinctions in network communication, we next examined how tract caliber– and myelin-weighted networks distribute communication across topological scales and how these differences relate to communication efficiency as a function of distance (Fig. 4). An extended spectral comparison is provided in Supplementary Material (Fig. S1).

**Figure 4.**
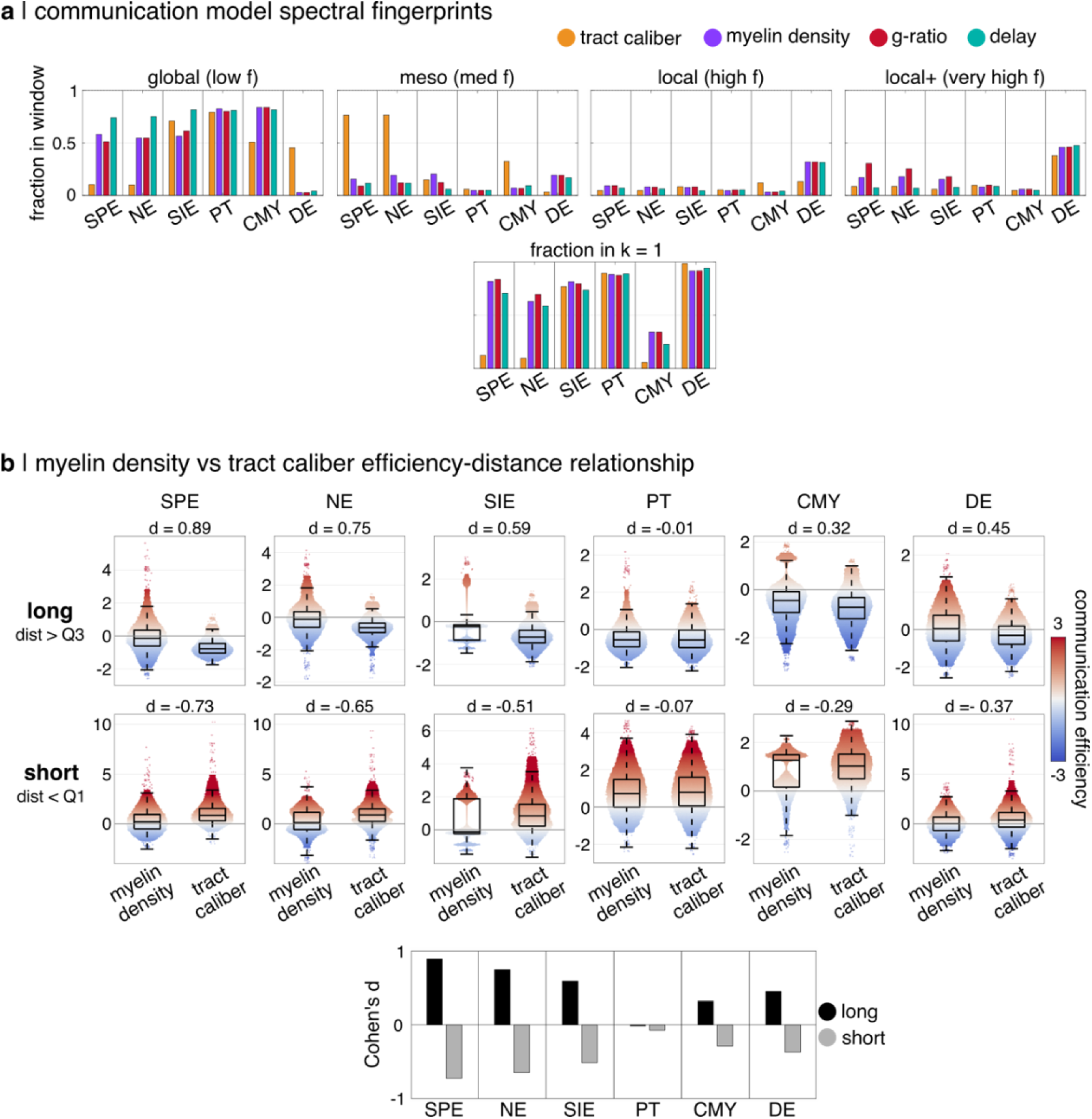
Functionally-relevant differences between tract caliber- and myelin-sensitive communication. (a) The fraction of spectral energy is shown for each topological window and separately for the 1^st^ eigenmode (k = 1). Colored bars represent structural networks grouped by communication model. Bar heights were normalized after separation of the 1^st^ eigenvector to prevent saturation of scale-dependent patterns. (**b**) Edgewise values of z-scored communication efficiency are compared between tract caliber and myelin density. The distribution of Euclidean distance between node pairs was used to group edges according to long (distance > 3^rd^ quartile) and short (distance < 1^st^ quartile). Cohen’s d was used to summarize the difference between distributions in each plot. Communication models included: shortest path efficiency (SPE), navigation efficiency (NE), search information efficiency (SIE), path transitivity (PT), communicability (CMY), and diffusion efficiency (DE).

#### Spectral fingerprints of communication models

We first used spectral fingerprinting to quantify how communication patterns for each model are distributed across eigenmodes of the underlying structural operator, from global to increasingly local scales (Fig. 4A). This approach provides a compact summary of the topological scales most strongly engaged by each communication model under different structural weightings.

Across the full range of eigenvectors excluding the 1^st^ global-mode (k 2-399), networks weighted by tract caliber and myelin exhibited differences in spectral energy distribution dependent on both communication strategy and topological scale. For routing-based models (e.g., SPE, NE), myelin-weighted communication (using myelin density, g-ratio, and delay) showed consistently higher spectral energy at more global modes, whereas tract caliber-weighted communication emphasized relatively finer meso-scale structure. Communicability followed the same qualitative pattern, though with somewhat attenuated differences between structural weightings. In contrast, diffusion efficiency exhibited the opposite trend: increased spectral energy from meso-scale to the smallest scale local+ modes for myelin-weighted communication, and a bias toward more global modes for tract caliber-weighted communication. These opposing patterns indicate that tract caliber and myelin weightings differentially redistribute communication across topological scales, and that the direction of this redistribution depends on the communication strategy.

Examination of the dominant eigenmode (k = 1) revealed pronounced differences in how structural weighting shapes communication at the most global scale. For routing-based models, myelin-weighted networks exhibited a substantially larger fraction of spectral energy at k = 1 (exceeding 0.7) compared to tract caliber-weighted networks, which showed minimal energy at this scale (< 0.1). Communicability again followed the same trend, with higher k = 1 energy for myelin-weighted networks (∼0.35) relative to tract caliber (< 0.1). Again, diffusion efficiency showed an inverted pattern at the global mode. Tract caliber-weighted DE was almost entirely dominated by the k = 1 mode (∼0.99), whereas myelin-weighted DE exhibited a modest reduction in k = 1 dominance (∼0.95), consistent with its redistribution of energy toward higher-order modes observed for k = 2–399. These findings demonstrate that differences in structural weighting are expressed most strongly at the largest scales of network organization, while also reshaping how communication engages mesoscale and local structure. Relative to tract caliber, myelin-weighting is associated with a redistribution of routing-based communication toward more global topological modes and of random-walker-based diffusive communication toward higher-order, mesoscale-to-local modes.

#### Distance-dependent communication efficiency

We next examined whether these topological differences correspond to variations in communication efficiency across spatial scales (Fig. 4B). Long-range connections were associated with consistently higher efficiency of myelin-weighted communication across all models except path transitivity (PT). Conversely, short-range connections were associated with higher efficiency for tract caliber-weighted communication for all models except PT. For both distance groups, the magnitude of Cohen’s d followed a U-shaped profile along the routing–diffusion spectrum of communication models: effect sizes were largest at either end of the spectrum and approached zero in the middle (SPE > NE > SIE > PT < CMY < DE). The largest effect sizes were observed for routing models (d = 0.89 and -0.73 for SPE of long and short distances, respectively). This pattern suggests that the functional consequences of structural weighting are most pronounced for communication strategies that strongly emphasize either direct routing or stochastic diffusion, with intermediate strategies showing reduced sensitivity.

Thus, networks weighted by tract caliber and myelin-sensitive metrics differ not only in their communication values, but also in the topological scales and spatial regimes over which communication is most efficient. Relative to tract caliber, myelin-weighted routing and communicability preferentially engage global structural modes, whereas myelin-weighted random-walker-based diffusion shifts communication toward meso- and local-scale structure. Despite this difference between models, myelin-weighted communication consistently supports more efficient long-range communication than tract caliber. Together, these results highlight a fundamental distinction in how tract caliber and myelin-sensitive weightings shape the balance between global integration and local processing in network communication.

### Global coupling of myelin-sensitive communication and functional connectivity

We next quantified the global contribution of myelin-weighted communication models to the prediction of functional connectivity across timescales using a nested regression framework (Fig. 5). Functional connectivity was examined across BOLD and electrophysiological frequency bands, allowing us to assess timescale-dependent structure–function coupling. For each communication model, the base regression model included Euclidean distance, tract caliber–weighted communication, and binary communication terms, thus capturing geometric and topology-driven contributions common across structural representations. The reported ΔR^2^ values therefore reflect the additional variance explained by incorporating myelin-weighted communication (myelin density, g-ratio and delay) beyond these baseline predictors. An assessment of communication-functional coupling using individual myelin-sensitive metrics is provided in Supplementary Material (Fig. S2-S6).

**Figure 5.**
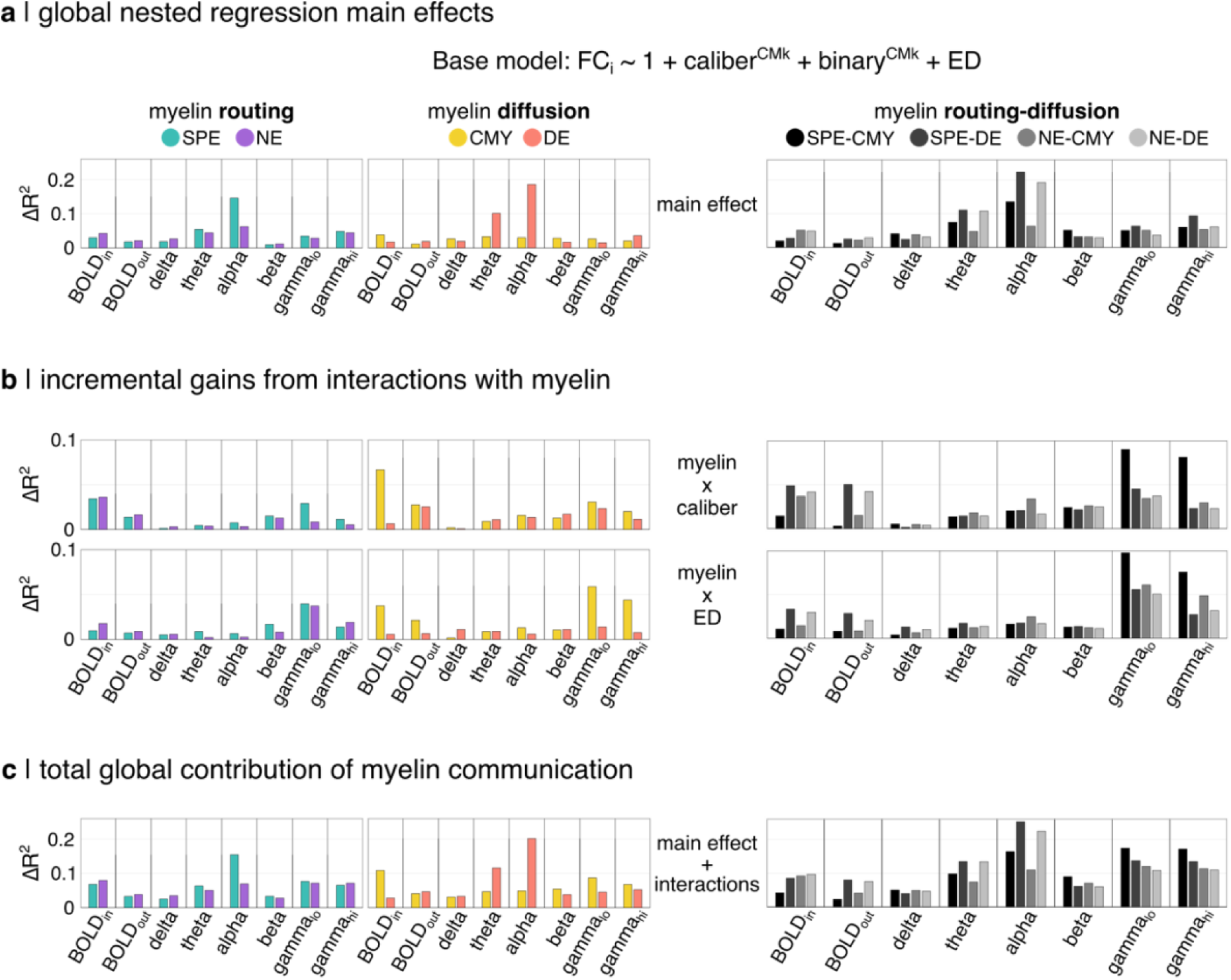
Global coupling of myelin-sensitive communication and functional connectivity (FC). Nested regression results are shown for (**a**) main effects, (**b**) incremental gains from interactions, and (**c**) the total contribution of myelin communication. The contribution of myelin-sensitive communication was computed relative to the base model: ΔR^2^ = full model – base model. The base regression model included tract caliber and binary communication measures, as well as Euclidean distance (ED). Nested F-tests were used to filter insignificant values (α = 0.05). Models were tested both individually and in routing-diffusion pairs. Interactions with myelin communication were examined for both tract caliber communication and Euclidean distance. Communication models included: shortest path efficiency (SPE), navigation efficiency (NE), communicability (CMY), and diffusion efficiency (DE).

#### Main effect of myelin-weighted communication

Across individual communication models (routing or diffusion), the largest gains in explained variance (ΔR²) were observed for alpha-band functional connectivity, particularly for routing-based models and diffusion efficiency (Fig. 5A). In contrast, communicability yielded substantially smaller ΔR² values, making peaks across timescales less visually distinct. Pairwise combinations of routing- and diffusion-based models produced higher ΔR² values than individual models for most functional connectivity datasets. The strongest coupling was again observed for alpha-band connectivity, followed by theta. Coupling with alpha was strongest for SPE-DE. These results suggest that alpha-band functional connectivity is most strongly constrained by global myelin-weighted communication processes, particularly when both routing and diffusion mechanisms are jointly considered.

#### Incremental gains from interactions with myelin-weighted communication

We next examined the incremental contribution from the interaction of caliber-weighted communication and Euclidean distance with myelin-weighted communication beyond the main effects accounted for above (Fig. 5B). Across models, the additional variance explained by interactions with myelin-sensitive communication tended to be larger at the edges of the functional timescale spectrum: larger ΔR² gains for BOLD and gamma-band connectivity than for theta and alpha. For both routing- and diffusion-based models, BOLD functional connectivity showed the largest gain from the interaction with tract caliber-weighted communication, whereas gamma-band connectivity exhibited the largest gain from the interaction with Euclidean distance. When routing and diffusion models were combined, the largest ΔR² gains from myelin-weighted communication interactions were observed for gamma-band connectivity across all interaction terms. The SPE–CMY combination showed particularly strong coupling. This pattern suggests that myelin-related modulation of communication contributes most strongly to functional connectivity at slower (BOLD) and faster (gamma) timescales, and that these effects depend on the specific structural feature with which myelin interacts.

#### Total global contribution of myelin-weighted communication

Finally, we assessed the total contribution of myelin-weighted communication by combining main effects and interaction terms (Fig. 5C). Across individual communication models (routing or diffusion), alpha-band connectivity remained a prominent peak but was less dominant then in the main effects condition, as BOLD and gamma-band connectivity now benefited from strong myelin interaction effects. Peaks for communicability became more apparent in both BOLD and gamma bands. When routing and diffusion models were combined, a similar redistribution was observed: alpha-band connectivity retained high ΔR² values, but gamma-band connectivity emerged as a close second. Thus, while alpha-band functional connectivity shows the strongest baseline coupling to myelin-weighted communication models, myelin-related interactions substantially enhance structure–function coupling at slower and faster timescales, reshaping the relative contribution of communication strategies across the frequency spectrum.

Taken together, these findings suggest that global structure–function coupling depends jointly on communication strategy, functional timescale, and interactions between myelin and other structural features. Alpha-band connectivity exhibits the strongest baseline association with myelin-weighted communication models, particularly shortest-path routing and random-walker-based diffusion, whereas BOLD and gamma-band connectivity derive a larger proportion of their explanatory power from myelin-dependent interactions. Thus, while myelin-sensitive modulation of communication plays an important role in linking structural networks to functional dynamics at the extremes of the temporal spectrum, the dominant contribution of myelin-weighted communication to functional connectivity prediction occurs at the mid-frequency timescales of alpha and theta.

### Regional heterogeneity of coupling between myelin-sensitive communication and functional connectivity

To determine whether coupling between myelin-weighted communication and functional connectivity varies across cortical systems and spatial scales, we measured the total myelin contribution (main effects + interactions) using nested regression at both the resting-state network (RSN) and node levels (Fig. 6–7). In Supplementary Material, RSN-level results are expanded to cover all functional connectivity datasets for ΔR2 (Fig. S7) and a permutation-based sensitivity index (Fig. S8). In addition, the relative magnitude of functional coupling is compared between myelin-sensitive and tract caliber-sensitive communication (Fig S9).

**Figure 6.**
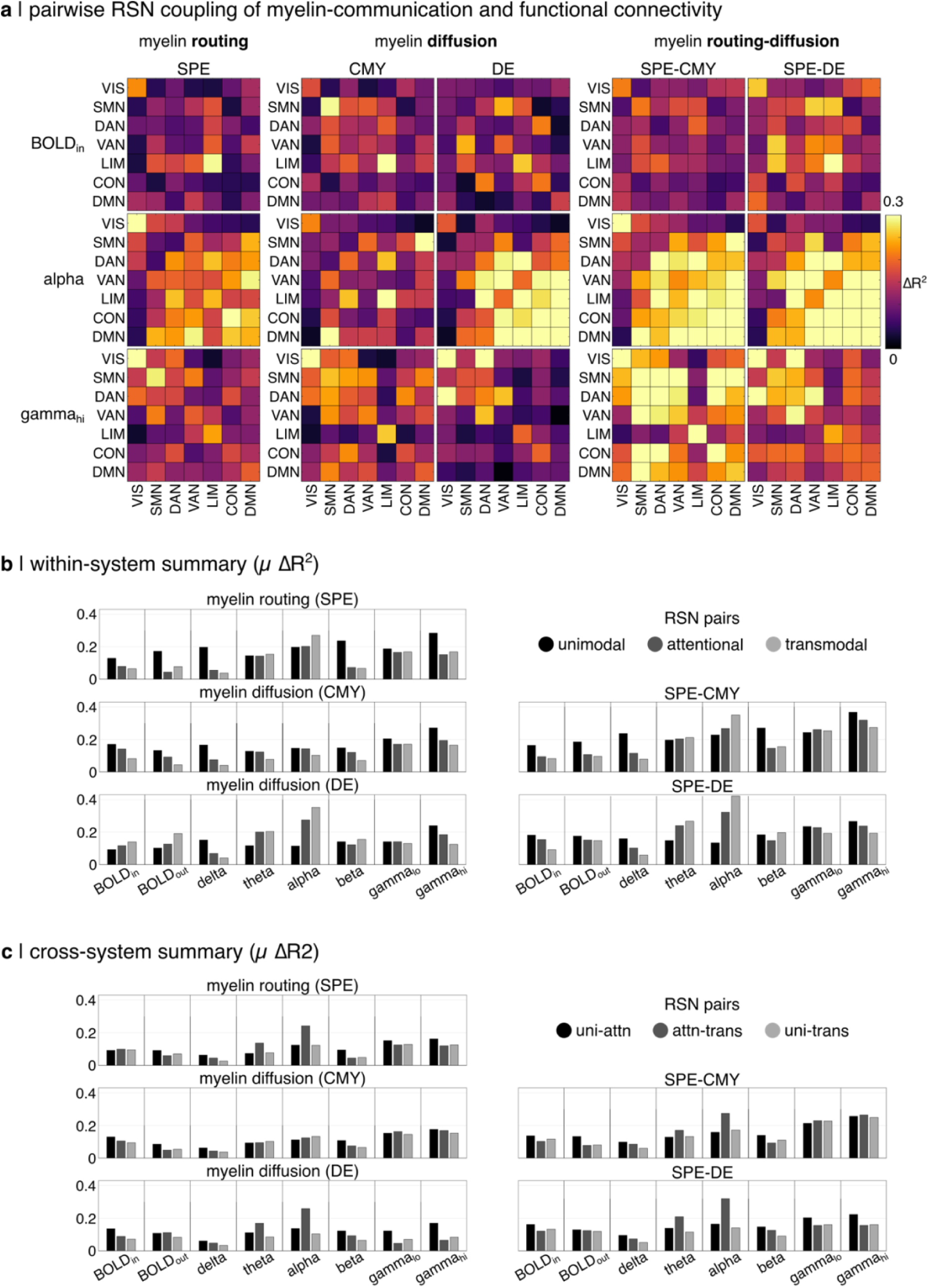
Network-level regional heterogeneity of coupling between myelin-sensitive communication and functional connectivity. (**a**) The total contribution of myelin communication (interactions + main effects) is shown for nested regression in pairwise combinations of resting-state networks. BOLD, alpha and gamma functional connectivity were selected to summarize the full spectrum of functional timescale (see Fig. S7 for more). The contribution of myelin-sensitive communication was computed relative to the base: ΔR^2^ = full model – base model. The base regression model included tract caliber and binary communication measures, as well as Euclidean distance (ED). Nested F-tests were used to filter insignificant values (α = 0.05). Models were tested both individually and in routing-diffusion pairs. Resting-state networks correspond to: visual (VIS), somatomotor (SMN), dorsal attention (DAN), salience ventral attention (VAN), limbic (LIM), fronto-parietal control (CON), default mode (DMN). (**b**) Within-system and (**c**) cross-system summaries of heatmaps are provided by averaging coupling values (µ ΔR^2^) according to canonical groupings of resting-state networks (unimodal: VIS, SMN; attentional: DAN, VAN; transmodal: CON, DMN). Communication models included: shortest path efficiency (SPE), communicability (CMY), and diffusion efficiency (DE).

**Figure 7.**
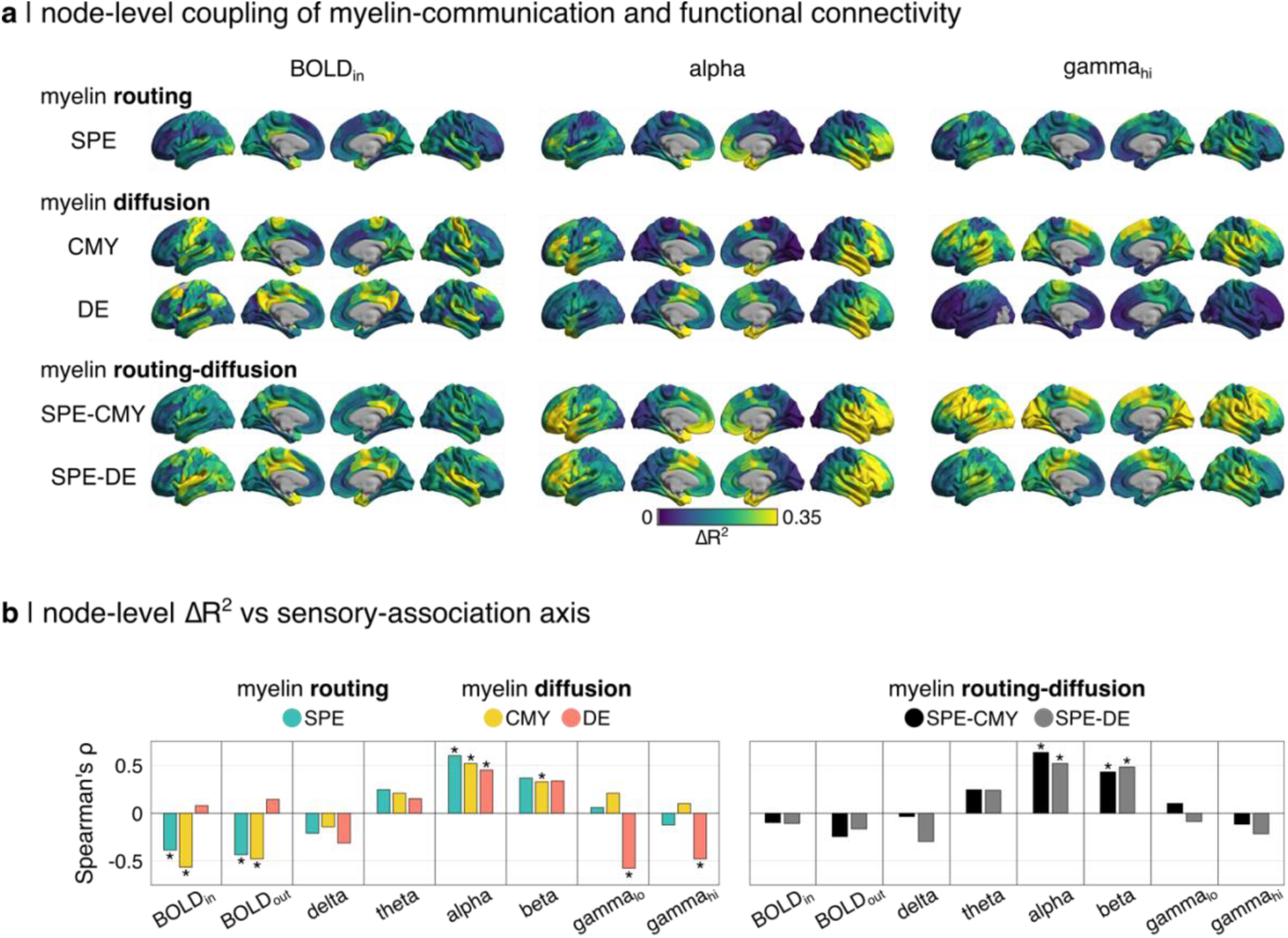
Node-level regional heterogeneity of coupling between myelin-sensitive communication and functional connectivity. (**a**) The total contribution of myelin communication (interactions + main effects) is shown for nested regression in individual nodes. BOLD, alpha and gamma functional connectivity were selected to summarize the full spectrum of functional timescale. The contribution of myelin-sensitive communication was computed relative to the base: ΔR^2^ = full model – base model. The base regression model included tract caliber and binary communication measures, as well as Euclidean distance (ED). Nested F-tests were used to filter insignificant values (α = 0.05). Models were tested both individually and in routing-diffusion pairs. (**b**) Spearman’s rank correlations of node-level coupling (ΔR^2^) with the sensory-association axis ordinal mapping are shown. Asterisks (*) indicate significance from hemispherically-constrained spin tests (n = 10,000, p_spin_ < 0.05). Communication models included: shortest path efficiency (SPE), communicability (CMY), and diffusion efficiency (DE).

#### Resting-state network organization of myelin–communication–FC coupling

At the RSN level, combining routing- and diffusion-based communication models increased both the magnitude and spatial extent of coupling with myelin communication across functional timescales. This effect was strongest for alpha- and gamma-band connectivity (Fig. 6A). The functional coupling of myelin-communication showed distinct spatial patterns across timescale.

For BOLD functional connectivity, coupling was generally weaker and more concentrated spatially than for higher-frequency bands. Within individual models, myelin-weighted shortest-path routing showed relatively stronger coupling within limbic and visual networks, whereas myelin-weighted diffusion (CMY and DE) showed higher coupling for network pairings involving somatomotor, attentional, and limbic systems. Combined routing–diffusion models amplified these effects, yielding peak coupling within limbic and visual networks and between somatomotor–attentional and somatomotor–limbic network pairs.

For alpha-band functional connectivity, coupling of individual myelin-weighted communication models was consistently strongest within attentional and heteromodal networks, with peak values observed both within these systems and between attentional–heteromodal network pairs. Diffusion efficiency showed the strongest alpha-band coupling overall. Combined routing–diffusion models further strengthened this pattern, producing robust coupling across attentional, heteromodal, and limbic systems.

Gamma-band functional connectivity showed a partial return toward unimodal systems. Individual models exhibited strongest coupling within sensory networks and between sensory and attentional systems. Communicability showed the highest coupling between the most network pairs. In combined models, SPE–CMY produced the strongest and most spatially extensive gamma-band coupling, including elevated within-network coupling across multiple RSNs.

Collectively, these results reveal a systematic shift in the spatial distribution of coupling with myelin-sensitive communication across functional timescales: from limbic and sensory systems in low-frequency BOLD, to attentional and heteromodal systems for mid-frequency alpha, and back toward sensory–attentional systems for high-frequency gamma coupling.

#### Node-level heterogeneity and cortical hierarchy

Node-level analyses refined these system-level patterns and revealed substantial spatial heterogeneity in coupling between myelin-sensitive communication and functional connectivity across the cortex (Fig. 7A). For BOLD connectivity, coupling was generally weaker and concentrated in posterior regions, including lateral occipital, temporal, and medial parietal cortex. Here, diffusion- and routing-based models showed partially distinct spatial profiles.

In contrast, alpha-band connectivity showed pronounced coupling in anterior cortical regions across all models. Peak coupling values were observed in lateral temporal and frontal cortex, with additional involvement of medial frontal and central regions. Coupling was consistently higher in anterior than posterior regions, and stronger laterally than medially. These effects were strongest and most spatially extensive for communicability but were present across all individual and combined models.

Gamma-band connectivity again exhibited a different spatial pattern. Communicability showed the strongest and most widespread coupling, with peaks spanning fronto-central, temporo-parietal, and occipital regions, whereas routing models exhibited more localized effects. Combined routing–diffusion models further amplified these patterns, with SPE–CMY producing broad, high-magnitude coupling across much of the cortex.

#### Alignment with the sensory–association axis

To assess whether spatial heterogeneity in myelin–communication–FC coupling aligns with cortical hierarchy, we examined nodewise coupling strength as a function of the sensory–association (S–A) axis (Fig. 7B). For individual models, BOLD connectivity showed significant negative correlations with the S–A axis for shortest paths and communicability, indicating stronger coupling in sensory regions. In contrast, alpha-band connectivity exhibited strong positive correlations with the S–A axis for all models, indicating preferential coupling in association cortex. Beta-band connectivity showed similar but weaker trends. For gamma-band connectivity, strong inverse correlations with the S–A axis were observed for diffusion efficiency only, and relationships were weak or absent for other models. For combined routing–diffusion models, alpha-band connectivity again showed robust positive correlations with the S–A axis, whereas relationships were weak or nonsignificant for BOLD and gamma bands. These results demonstrate a strong and frequency-specific alignment between myelin–communication–FC coupling and cortical hierarchy, with alpha-band connectivity showing the most consistent and robust association with the sensory–association axis.

Overall, coupling between myelin-sensitive communication and functional connectivity is highly heterogeneous across cortical systems, spatial scales, and functional timescales. Alpha-band connectivity exhibits the strongest and most spatially specific coupling, particularly within attentional and heteromodal networks and along the cortical hierarchy. In contrast, BOLD and gamma-band connectivity show stronger coupling in sensory and limbic systems, with gamma-band effects being most pronounced for communicability-based models. These findings suggest that myelin-sensitive modulation of network communication engages distinct cortical systems depending on functional timescale, with intermediate frequencies preferentially coupling to association cortex and extremes of the temporal spectrum engaging more sensory and limbic circuitry.

### Sensitivity analyses

We assessed the robustness of the main findings to analytic choices and dataset composition. Using the same participants and processing pipeline, results were broadly consistent at a lower parcellation resolution (Schaefer-200; Fig. S10), although the interaction of tract caliber and myelin communication made weaker contributions to functional connectivity prediction. We also observed a similar pattern in an independent multimodal dataset (10.17605/OSF.IO/J532R), in which myelin-communication–FC coupling was again strongest for shortest paths and diffusion efficiency in the alpha band (Fig. S11). Although this dataset did not include the tract-specific g-ratio network available in the primary sample, it indexed white matter myelin using the R_1_ relaxation rate (1/T_1_). These analyses suggest that the main effects are robust across datasets, parcellation scales, and myelin-sensitive metrics.

## Discussion

In this work, we demonstrate that incorporating myelin-sensitive information into communication models selectively strengthens structure-function coupling in a context-dependent manner shaped by communication strategy, cortical hierarchy, and functional timescale.

Among the functional signals examined, alpha-band connectivity exhibited the strongest coupling to myelin-sensitive communication models. Although present at the global level, this effect was amplified regionally, revealing heterogeneous coupling across resting-state networks and cortical regions. This suggests that alpha-band dynamics may occupy a privileged regime in which the structural constraints imposed by myelinated pathways are most directly expressed. In contrast, coupling in BOLD and higher-frequency bands was weaker, more spatially diffuse, or more contingent on interactions with other structural features, consistent with these signals reflecting slower integrative processes or more local dynamics.

Finally, routing- and diffusion-based strategies engaged distinct aspects of network topology and emphasized different cortical systems. Routing models preferentially highlighted mesoscale organization and rank-ordered pathways, whereas diffusion models were more sensitive to global topology and spatial embedding. These effects were further modulated by structural weighting (myelin-sensitive metrics vs. tract caliber), underscoring that white matter structure supports multiple, feature-dependent communication regimes that couple to functional connectivity in distinct ways.

### Myelin, topology, and communication strategy

The frequency-specific coupling of myelin-sensitive communication and functional connectivity can be understood by considering how communication models engage network topology at distinct scales and how structural features bias these engagements. We find a systematic divergence between routing- and diffusion-based strategies, wherein tract myelin and caliber differentially emphasize global and mesoscale organization.

In the spectral domain, low-order eigenvectors capture broad, integrative modes of network organization, whereas intermediate components reflect mesoscale patterns such as modular structure and systems-level segregation^32–34^. Under routing-like communication strategies (SPE, NE), myelin-sensitive weighting shifted communication toward globally integrative topological modes. Spectral fingerprinting revealed that myelin-weighted routing concentrated energy in low-order components, including the leading eigenmode, indicating stronger engagement of integrative network structure. In contrast, caliber-weighted routing preserved greater spectral energy at intermediate components, reflecting stronger engagement of mesoscale topology. Thus, while tract caliber-weighted routing sharpens modular differentiation, myelin-sensitive routing coordinates inter-modular communication across a globally integrative backbone.

This global bias under myelin-sensitive routing was further supported by edge-length analyses. Myelin-weighted communication efficiency increased preferentially for longer connections, especially for routing models. Long-range edges disproportionately contribute to network integration^35–37^, and their selective enhancement under myelin weighting links microstructural variation to low-order topological dominance. In contrast, caliber-weighted routing models showed increased communication efficiency for short-range connections, in agreement with their greater sensitivity to local and mesoscale structure.

Differences in community organization provide convergent evidence for this interpretation. Myelin-weighted networks formed communities that were spatially diffuse, consistent with integrative, system-spanning organization. In contrast, caliber-weighted networks exhibited more spatially localized communities, reflecting stronger segregation and mesoscale differentiation. These community patterns align with the spectral and edge-length results in pointing to a bias toward integration under myelin-weighted routing and segregation under caliber-weighted routing.

Diffusion-based strategies exhibited a distinct profile. Diffusion was dominated by global topology, with spectral energy concentrated in low-order components regardless of structural weighting. Within this globally constrained regime, myelin and caliber exerted opposing secondary effects: caliber-weighted diffusion further reinforced low-order dominance, whereas myelin-weighted diffusion retained relatively greater intermediate-scale contributions. These results suggest that, when communication is distributed across many paths, global embedding constrains signaling across structural features, and microstructural differences modulate residual topological structure.

Correlations between tract caliber- and myelin-sensitive communication further clarify how structural features interact with communication strategy. Under routing models, myelin- and caliber-based communication were inversely correlated for connected edges and uncorrelated for disconnected edges, indicating that these features prioritize competing rather than redundant routing pathways. Under diffusion models, correlations were positive for disconnected edges and weak for connected edges, suggesting convergence toward shared global embedding constraints when communication is distributed. Together, these findings show that the influence of white matter microstructure on inter-regional signaling depends on both the structural feature emphasized and the assumed communication strategy.

### Temporal scale, cortical hierarchy, and regionally heterogeneous coupling

The prominence of alpha-band functional connectivity in myelin–communication–function coupling underscores the importance of temporal scale in determining how structural constraints are expressed. Alpha oscillations^38^ operate at an intermediate timescale that is well positioned to interact with large-scale white matter architecture, providing a bridge between structural topology and functional dynamics^2,39^. One plausible explanation for this privileged role lies in timescale matching between neural dynamics and inter-regional communication delays^40,41^. Conduction delays along myelinated white matter pathways fall within a range compatible with the period of alpha oscillations^6,11,42^, allowing distributed regions to interact coherently without excessive phase dispersion. In this sense, alpha rhythms may resonate with the temporal constraints imposed by network structure, allowing myelin-sensitive structural features to exert a stronger influence on functional coupling. This interpretation is consistent with evidence that large-scale synchronization is most effective when oscillatory periods align with communication delays^41,43^, with the importance of alpha in delay-constrained neural mass models^44^, and with the established role of alpha rhythms in coordinating distributed cognitive processes^38,45^.

Coupling patterns were regionally heterogeneous at both the systems and node levels, supporting the idea that structure–function relationships depend on both the computational role of a region and the timescale of the dynamics under consideration. The expression of alpha-band coupling aligned systematically with the sensory–association axis: weaker coupling in primary sensory cortex and stronger coupling in higher-order association regions. This pattern is aligned with both hierarchical differences in intrinsic timescales^46,47^ and integrative demands across the cortex^48,49^. Association areas operate on slower, more integrative dynamics and rely more heavily on long-range coordination, whereas sensory regions are dominated by faster, locally driven processes. The alignment of alpha-band coupling with this hierarchy suggests that myelin-sensitive communication is most influential when both temporal integration and distributed coordination are required.

Attentional networks also exhibited strong coupling between myelin-sensitive communication and functional connectivity. Although present at other frequencies, this coupling was particularly strong in the alpha band, which is conceptually compatible with the putative role of attentional systems as flexible coordinators of distributed processing. Positioned between unimodal sensory and heteromodal association systems^48^, attentional networks support flexible reconfiguration across tasks and states^50,51^. Their reliance on rapid, selective, long-range coordination may render them especially sensitive to microstructural constraints on inter-regional signaling. Together, these findings indicate that intermediate temporal scales preferentially engage distributed attentional and association systems in which myelin-sensitive communication exerts maximal influence.

At the frequency extremes, a different pattern emerged. In BOLD-derived functional connectivity, structure–function coupling was more spatially diffuse and more strongly shaped by interactions with tract caliber and Euclidean distance, particularly under communicability-based models. This profile is consistent with the temporally integrated and spatially smooth nature of haemodynamic signals, which increases sensitivity to aggregate communication capacity and spatial embedding^1,2,52^. In gamma-band connectivity, coupling was more localized and dominated by interactions with Euclidean distance, reflecting a greater reliance on short-range circuitry and temporally precise synchronization^39,53^. Across both extremes, coupling patterns shifted toward sensory systems, suggesting that when functional dynamics are either strongly temporally integrated (BOLD) or highly localized (gamma), structure–function relationships are more heavily governed by spatial proximity and short-range architecture than by large-scale myelin organization.

Collectively, these findings demonstrate that the influence of white matter microstructure on functional connectivity is jointly constrained by temporal scale and cortical hierarchy. Alpha-band activity occupies a regime in which realistic conduction delays, large-scale topology, and hierarchical organization align to amplify the expression of myelin-sensitive communication. In contrast, slower or faster signals reflect different balances between spatial embedding, short-range structure, and interactions among multiple structural features. This regime-dependent pattern underscores that structure–function coupling is neither spatially nor temporally uniform, but emerges from the interaction between network topology, microstructural weighting, and the dynamical scale of neural activity^17^.

### Mechanistic interpretations of myelin-sensitive communication

While the present analyses cannot adjudicate specific cellular or biophysical mechanisms, the convergence of topology-, frequency-, and region-specific effects constrains how myelin-related microstructure may shape large-scale functional communication. Below, we outline several non–mutually exclusive mechanistic interpretations consistent with these constraints.

A natural starting point is the classical role of myelin in modulating conduction velocity^7^ and inter-regional signaling delays^11^. Although the coupling between myelin-sensitive communication and alpha-band connectivity aligns with the timescale-matching framework described above, conduction delay alone is unlikely to explain the observed patterns. If myelin influenced functional coupling solely through conduction velocity, one would expect substantial redundancy between tract caliber– and myelin-weighted communication, given the joint dependence of delay on axon diameter and myelination^54,55^. Instead, these features exhibited dissociable effects across communication strategies and topological scales, suggesting that myelin contributes beyond simple modulation of signal delay.

An alternative interpretation is that myelin acts as a permissive or stabilizing factor for long-range communication, particularly in distributed association and attentional systems. Long-distance white matter pathways are metabolically costly^37,56^ and vulnerable^57,58^, and their sustained functional engagement may depend on trophic and maintenance support from oligodendrocytes^9,59^. From this perspective, myelin-sensitive measures may index not only transmission efficiency, but also the reliability and long-term viability of inter-regional signaling pathways. This framing helps explain why myelin-weighted communication emphasized globally integrative topology and was strongest in systems supporting flexible long-range coordination.

A further possibility is that myelin influences functional connectivity through energetic constraints and their interaction with population-level dynamics. Myelination reduces the metabolic cost of action potential propagation^56^, which may preferentially support sustained large-scale coordination^37^. Alpha-band connectivity, relative to higher-frequency rhythms, reflects broader population synchrony over longer integration windows and lower average firing rates^38^. The association between myelin-sensitive communication and alpha coupling may therefore reflect a regime in which energetic efficiency stabilizes distributed oscillatory coordination. This does not imply that myelin determines oscillatory frequency per se, but rather that certain frequency regimes may be more sensitive to microstructural constraints on energy-efficient long-range signaling.

In sum, these considerations support a regime-dependent view of myelin-sensitive communication in which multiple mechanisms jointly shape structure–function coupling. Tract caliber and myelin may jointly shape conduction delays, while myelin further modulates the reliability, energetic efficiency, and functional viability of inter-regional signaling. Communication strategy then determines which of these properties becomes most relevant. The central insight emerging from our results is therefore not the identification of a single mechanistic pathway, but the demonstration that myelin-related microstructural features exert their strongest influence on functional connectivity under specific combinations of temporal scale, network topology, and regional specialization. This perspective provides a biologically grounded framework for interpreting myelin-sensitive communication effects.

### Connection to prior work

A large body of work has examined structure-function coupling(e.g., (Baum et al., 2020; Demirtaş et al., 2019; Liu et al., 2023; Murray et al., 2018; Popp et al., 2024; Preti & Van De Ville, 2019; Vázquez-Rodríguez et al., 2019; Wang et al., 2019; Zamani Esfahlani et al., 2022). We previously advanced this literature by revealing regional- and frequency-dependent coupling between white matter myelin and functional connectivity, including the first demonstration of a modulatory role of myelin in structure–function coupling^17^. However, this modeling approach utilized the edge weights directly and thus was unable to account for most functional interactions. Here, we integrate biologically weighted structural connectomes, communication modeling, and functional connectivity at different temporal frequencies within a unified framework. By modeling higher-order myelin-sensitive communication between indirectly connected regions, we show that strong structure–function coupling in association cortex persists in the alpha band, contrasting with reports of gradual decoupling along the sensory–association axis derived primarily from BOLD fMRI. These findings suggest that hierarchical gradients in structure–function coupling are modality-, timescale-, and microstructure-dependent, rather than universal features of cortical organization.

This work also advances network communication modeling beyond conventional streamline-count or density-based representations^18,31^. By embedding myelin-sensitive measures directly into communication frameworks, we show that structural weighting does not merely rescale communication estimates but qualitatively shifts which topological regimes are engaged and how coupling is expressed across cortical systems and frequencies. These findings position microstructural weighting as a critical dimension in communication modeling, moving these approaches toward a more mechanistically interpretable account of large-scale brain dynamics.

### Limitations and future directions

Several limitations delineate the interpretational scope of the present findings. The analyses are inherently correlational and model-based and thus cannot adjudicate specific cellular or biophysical mechanisms. Moreover, communication models provide abstractions of how signals may traverse the structural connectome rather than direct measurements of neural transmission. Future work combining connectomic modeling with experimental manipulations, more detailed models^64,68^, cohorts exhibiting greater white matter variability^69–72^, or alternate communication^19,73^ or connectivity^74,75^ frameworks will be required to more directly probe causal mechanisms and refine mechanistic inference^76^.

The structural features considered here are indirect biological proxies. Tract caliber estimates a bundle-level property, and myelin-sensitive measures (MTsat, g-ratio, delay) provide complementary but incomplete indices of white matter microstructure. Neither class captures axonal heterogeneity^28^, synaptic efficacy^77^, neuromodulatory influences^78^, activity-dependent myelin plasticity^8^, or intra-cortical myelin^79,80^, all of which may shape functional communication. Integrating richer microstructural imaging, longitudinal designs, and molecular or metabolic markers may further clarify how white matter architecture supports large-scale functional organization.

The MEG data analyzed here were obtained from an out-of-sample cohort, precluding subject-level structure–function analyses across modalities. Future work combining structural MRI, BOLD fMRI, and electrophysiology within the same individuals will be important for assessing inter-individual variability^81^ and determining whether the regime-dependent effects observed here extend to subject-specific predictions.

Finally, the present analyses focus on static, resting-state functional connectivity in healthy adults. Communication regimes and their sensitivity to microstructural constraints may differ across tasks^82^, dynamic functional states^83^, development^63^, aging^84^, or neurological and psychiatric conditions^85^. Extending this framework to task-based, dynamic, longitudinal, and clinical functional data represents an important direction for future research.

## Conclusion

Our findings demonstrate that white matter microstructure shapes functional connectivity in a context-dependent manner. By integrating myelin-sensitive structural measures with network communication models and frequency-resolved functional connectivity, we show that microstructural properties selectively constrain inter-regional coupling in oscillatory regimes and cortical systems that support distributed, integrative coordination. In particular, alpha-band functional connectivity emerges as a privileged regime in which realistic communication delays, energetic efficiency, and large-scale topology align to amplify the expression of myelin-sensitive communication. Together, these results advance a multiscale and biologically grounded account of structure–function coupling and underscore the importance of incorporating microstructural specificity into models of large-scale brain communication.

## Materials and Methods

### Data acquisition

This study was approved by the Research Ethics Board of McGill University, and all participants provided written informed consent. Multi-modal MRI data were collected in a cohort of healthy volunteers (n = 30; 14 men; 29±6 years of age) on a 3 tesla Siemens Magnetom Prisma-Fit scanner equipped with a 64-channel head coil. All sequences used the Siemens *Prescan Normalize* option to generate maps with minimal receive B_1_-field bias. The protocol was as follows:

- T_1_-weighted (T_1_w) anatomical: 3D magnetization-prepared rapid gradient-echo sequence (MP-RAGE; 1.0 mm isotropic; TE/TI/TR = 2.98/900/2300 ms; FA = 9°; iPAT = 2).
- T_1_ relaxometry: compressed sensing 3D-MP2RAGE^86,87^ sequence (1.0 mm isotropic; TE/TI_1_/TI_2_/TR = 2.66/940/2830/5000 ms; FA_1_/FA_2_ = 4°/5°; TF = 175; undersampling factor = 4.6.
- Resting-state fMRI: multiband accelerated 2D-BOLD gradient echo echo-planar sequence (2.6 mm isotropic; TE/TR = 30/746 ms; MB factor = 6; FA = 50°). Two spin-echo images with AP and PA phase encoding were additionally acquired for distortion correction (2.6 mm isotropic; TE/TR = 60.6/6160 ms; FA = 90°).
- Magnetization transfer-weighted 3D gradient-echo sequence (1.0 mm isotropic; TE/TR = 2.76/27 ms; FA = 6°; iPAT = 2; partial Fourier = 6/8; MT pulse parameters: 12-ms Gaussian pulse, 2kHz off-resonance, B1_rms_ = 3.26μT.
- Multi-shell diffusion-weighted imaging (DWI): 2D pulsed gradient spin-echo echo-planar imaging sequence composed of three shells with b-values 300, 1000 and 2000 s/mm^2^ and diffusion directions 10, 30, and 64, respectively (2.6 mm isotropic; TE/TR = 57/3000 ms; iPAT = 3; partial Fourier = 6/8). Two b = 0 images with AP and PA phase encoding were additionally acquired to facilitate distortion correction.
- A B ^+^ map acquired using the presaturated turboFLASH sequence^88^ (2.5 x 2.5 x 3mm^3^; TE/TR = 2.22e^-3^/20 s; FA = 8°) for ΔB_1_ ^+^ correction of MP2RAGE T1 & MTsat maps.

### Data processing

A custom version of the multi-modal processing pipeline *micapipe* (v0.1.5)^89^ was used in the processing of all diffusion, anatomical, and functional images. This pipeline makes use of multiple software packages including: *AFNI*^90,91^, *FSL*^92–94^, *ANTs*(B. Avants et al., 2009; Brian B. Avants et al., 2014), *FreeSurfer*^97–102^, and *MRtrix3*^103,104^.

#### T1w anatomical MRI (MPRAGE)

T_1_w images were deobliqued, reoriented to standard neuroscience orientation (LPI), N4 bias field corrected^105^, intensity normalized, and skull stripped^106^. Model-based segmentation of subcortical structures was performed with FSL FIRST^107^ and tissue types were classified using FSL FAST^108^. A five-tissue-type image segmentation was generated for anatomically constrained tractography^109^. Cortical surface segmentations were generated with the *FreeSurfer* (v7.2) *recon-all* pipeline with manual edits to remove dura mater.

#### Resting-state fMRI

After dropping the first 5 TRs from the main scan, all fMRI images (including field maps) were reoriented to standard neuroscience orientation (LPI) and motion corrected within-scan by registering all volumes to their respective temporal mean. Framewise displacement was computed for all main scan volumes, and a threshold (3^rd^ quartile + 1.5 * Interquartile Range) was used to identify volumes with large motion artifacts. The main-phase and reverse-phase field maps were used to correct the main fMRI scan for geometric distortions due to magnetic field inhomogeneity^110,111^. All subsequent processing involves only the main fMRI scan. Brain extraction^112^ was performed, followed by registrations to native FreeSurfer space with *bbregister*^113^ and with the T_1_w image using a multi-stage method (rigid, affine, SyN) implemented in ANTs^114^.

The fMRI data was denoised by applying a high-pass filter at a cutoff frequency of 0.01 Hz followed by Independent Component Analysis with FSL MELODIC(C.F. Beckmann & Smith, 2004; Christian F. Beckmann et al., 2005). All independent components (n = 3,825) were manually labeled by 2 independent raters, and components labeled as “noise” by both raters were removed (n = 2761). The denoised data was then used to calculate tissue-specific signals for gray matter, white matter, and cerebrospinal fluid tissue classes, before being mapped to native surface space and smoothed (Gaussian, FWHM = 6 mm). Linear regression was applied to the surface-based time series to remove nuisance signal resulting from motion, cerebrospinal fluid or white matter tissue.

#### Multi-shell DWI

The DWI data including field maps (b = 0 s/mm^2^ volumes with reverse phase encoding) were denoised^117–119^, then corrected for Gibbs ringing^120^. The field maps were then used to correct for susceptibility distortion, head motion, and eddy currents via FSL’s TOPUP^110^ and eddy^121^. This was followed by B_1_ ^+^ bias-field correction^105^. The corrected DWI image was upsampled to match the T1w resolution (1 mm isotropic), and brain extraction was performed^112^. These preprocessed data were used to estimate multi-shell and multi-tissue response functions^122–124^, which informed the estimation of fiber orientation distributions (FOD) via spherical deconvolution^125^. All FODs were intensity normalized^126^. A multi-stage (rigid, affine, SyN) registration^114^ was computed with the T_1_w image to allow co-registration of anatomical images to DWI space.

#### T_1_ relaxometry and magnetization transfer saturation (MTsat) maps

A dictionary matching approach incorporating the B ^+^ map was applied to generate S_0_ and T_1_ maps from the MP2RAGE inversion 1, inversion 2 and UNI images. This produced ΔB_1_ ^+^ corrected S_0_ & T_1_ maps. An affine transformation (ANTs) was used to register the MP2RAGE denoised UNI image, the S_0_ and T_1_ maps, and the MT-weighted image to the T_1_w image. MTsat maps were computed using^127^:

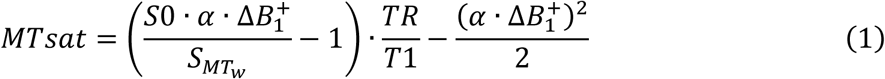

where *α* is the excitation flip angle in the MT-weighted sequence, *S*_*MTw*_ is the MT-weighted signal, and TR is the repetition time of the MT-weighted sequence. A gain factor of 2.5 was applied to the S_0_ maps to match the receiver gain of the MT-weighted gradient echo sequence for computing MTsat. ΔB_1_ ^+^ was corrected using a model-based approach^128^.

#### Voxel-wise g-ratio mapping

Voxel-wise myelin and axonal volume fractions were first derived from MTsat and DWI data, respectively. Intra-cellular (ICVF) and isotropic (ISOVF) volume fractions were estimated using the NODDI model within the AMICO framework^129^. MTsat maps were converted to myelin volume fraction (MVF) following calibration in the splenium of the corpus callosum, where a g-ratio of 0.7 was assumed^130,131^. A calibration factor (α_calib_) was computed by first pooling all voxels with FA > 0.8 within the splenium across the entire cohort, then averaging and calculating MVF as:

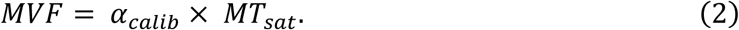

The voxel-wise axonal volume fraction (AVF) was then obtained from the relationship

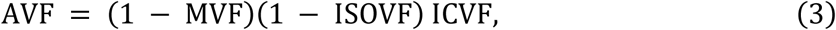

and the corresponding g-ratio map was calculated as,

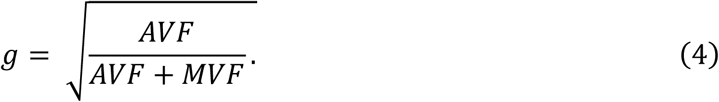

### Structural connectivity network reconstruction

#### Tractography, filtering and tract-specific volume estimation

Anatomically constrained^109^ tractography was run on the normalized white matter FOD with the probabilistic algorithm iFOD2^132^ and a dynamic seeding strategy^133^ to generate tractograms of 3 million streamlines. In a multi-stage process, these tractograms were filtered and both axonal and myelin volumes were computed for all surviving streamlines:

1. Any erroneous streamlines that failed to connect two gray matter ROIs were manually removed to ensure they did not bias estimates in subsequent steps.
2. COMMIT v2.1 ^26,27^ (convex optimization modeling for microstructure informed tractography) was run on the stage-1 filtered tractograms. The model was optimized to the DWI image using a Stick-Zeppelin-Ball forward model with diffusivities Ð_∥_ = 1.7e^-3^, Ð_⊥_ = 0.51e^-3^ and Ð_*iso*_ = 1.7e^-3^, 3.0e^-3^ mm^2^/s (gray matter and CSF). Estimates of the signal fraction or ICVF of each streamline were extracted from the COMMIT model. Streamlines with weight < 1e^-12^ were filtered as they did not contribute to the global DWI signal. Assuming a splenium g-ratio of 0.7, the ICVF map generated by COMMIT was used to recalibrate the MTsat-derived MVF map.
3. The stage-2 filtered tractograms were optimized to the MVF map with a myelin-based COMMIT extension^134^ yielding a signal fraction for each streamline which corresponded to a myelin cross-sectional area. Streamlines with weight < 1e^-12^ were filtered as they did not contribute to the global MVF signal. Surviving streamline weights were multiplied by streamline length to yield the myelin volume (MV) of each tract.
4. Because myelin exhibits negligible signal in DWI due to rapid T₂ decay, the intra-axonal volumes from step 2 overestimate the true axonal contribution. To correct for this, the diffusion signal was scaled by (1 − MVF) at the voxel level and normalized by the non-diffusion-weighted signal (b₀). These adjusted data were then upsampled to 1 mm isotropic resolution and re-fit with COMMIT with the “doNormalizeSignal” option disabled. The streamline weights output by COMMIT in this configuration represent their true axonal cross-sectional areas. During this stage, a minimal number of streamlines with weight < 1e^-12^ were filtered. Finally, true axonal volumes (AV) were derived by multiplying the surviving streamline weights by their length.

#### Weighted structural connectomes

For each subject, the same filtered tractogram was used for all structural connectomes, and the Schaefer-400 cortical atlas^25^ was used to define network nodes in all primary analyses. The edge *length* connectome was computed as the mean length of streamlines for each node pair. The edge *caliber* connectome was computed by summing the final COMMIT weights (signal fractions) of all remaining streamlines connecting each node pair normalized by total node volume for each pair. The edge *MTsat-myelin* connectome (myelin density) was computed by applying tractometry^135^ to the MTsat image: the median MTsat value was computed along each streamline, followed by the average across streamlines for each node pair. For the edge *g-ratio-myelin* connectome, MV and AV connectivity matrices were generated by summing the respective streamline volumes across each node pair, and tract-specific g-ratio was computed element-wise as

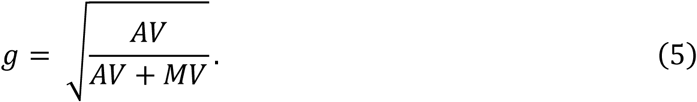

#### Group-consensus structural connectomes

We developed a group averaging method through rigorous testing to maximally preserve subject-level features. This was carried out in a two-stage process. (1) Consistency-based filtering^136^ targeted to cross-subject variance in number of streamlines edge weights was used to filter all subject-level structural connectomes to a threshold of 30% (the final density if averaging was performed at this stage). (2) Distance-dependent thresholding^137^ was then applied to generate group-representative structural connectomes which preserved subject-level within- and between-hemisphere edge length distributions. Final group-level connection density was 23%, which was approximately the median subject-level density.

#### Derivation of the delay connectome

To estimate tract conduction velocity and delay from the g-ratio-weighted connectome, we applied the Rushton biophysical model^54^, which predicts conduction velocity *v* as proportional to axon diameter *d* divided by the g-ratio *g*:

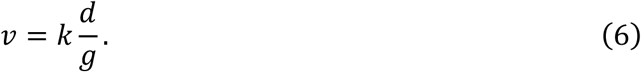

In the absence of reliable *in vivo* estimates of axon diameter, a constant *d* = 2.5 µm was assumed, with a proportionality constant *k* = 6.0 m s⁻¹ µm⁻¹ following prior work^138^. The resulting velocity matrix was combined with the edge length matrix (L) to compute conduction delays as

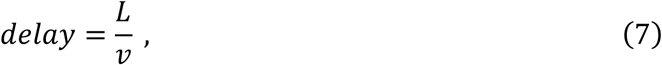

yielding an edge *delay* connectome expressed in milliseconds. Finally, each delay edge was inverted to obtain a 1/delay connectome, representing the effective communication rate between nodes. This was done to provide a complementary, weight-like representation of conduction efficiency. Whereas the delay connectome emphasizes transmission cost in temporal units (ms), the 1/delay formulation expresses the same information as an effective connection strength, enabling direct integration with standard weighted topology and communication model analyses.

### Functional connectivity network reconstruction

Functional connectivity was computed as the zero-lag Pearson cross-correlation of fMRI time series. These values were Fisher z-transformed and averaged across subjects yielding a single in-sample-fMRI functional connectome in the Schaefer-400 parcellation.

#### HCP data

Fully-processed^24^ fMRI and MEG functional connectivity matrices were obtained at the group-level in the Schaefer-400 parcellation. These connectomes were derived from resting-state data acquired in healthy young adults (n = 33; age range 22 to 35 years) as part of the Human Connectome Project (HCP; S900 release^139^). The MEG data corresponded to 6 minutes of scanning (sampling rate = 2,034.5 Hz; anti-aliasing low-pass filter = 400 Hz), and the fMRI data were comprised of 4 scans of 15 minutes across 2 sessions (TR = 720 ms; 3 Tesla; 2 mm isotropic). Static functional connectivity was computed as (1) the Pearson cross-correlation of fMRI time series and (2) the amplitude envelope correlation^140^ of MEG time series data. An orthogonalization process was then applied to correct for spatial leakage in the MEG functional connectomes^141^. Group-averaging yielded one fMRI and six MEG functional connectomes corresponding to the canonical electrophysiological frequency bands: delta (δ: 2-4 Hz), theta (θ: 5-7 Hz), alpha (α: 8-12 Hz), beta (β: 15-29 Hz), low gamma (γ_lo_: 30-59 Hz), and high gamma (γ_hi_: 60-90Hz). See^24^ for full processing details.

### Community detection and cross-partition comparison

For each weighted, undirected, structural connectivity matrix A, we (i) enforced symmetry (A ⇓ (A + A^T^) / 2), (ii) z-scored nonzero weights and applied a small positive shift such that all nonzero entries were > 0 (to avoid sign issues in weighted modularity), and (iii) set missing values to 0. No density thresholding was applied at any resolution step; the weighted graph was fixed across resolution sweeps.

#### Resolution parameter sweep

Communities were detected by maximizing weighted modularity with a tunable resolution parameter γ:

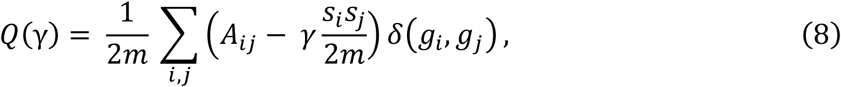

where *s*_*i*_ = ∑_*j*_ *A*_*ij*_, *g*_*i*_ is the community to which node *i* is assigned, the *δ*-function *δ*(*u*,*v*) is 1 if *u* = *v* and 0 otherwise, and 2*m* = ∑_*ij*_ *A*_*ij*_. We used the Louvain algorithm and varied γ over a log-spaced range to probe mesoscopic scales. This was implemented in a two-pass sweep of γ values. (1) The initial coarse sweep sampled *γ* ∈ [0.1, 5] (log-spaced, 100 samples) and performed a single optimization per γ value. We identified the first γ yielding > 2 communities (γ*_u_*) and the last γ yielding < N / 2 + 1 communities (γ*_v_*) as the bounds of the stable meso-scale regime. (2) The refined sweep sampled *γ* ∈ [*γ*_*u*_, *γ*_*v*_] (log-spaced, 200 samples) and performed 100 independent Louvain optimizations (repetitions) for each γ.

#### Identifying optimal partitions

For each γ, we quantified repeatability or similarity of partitions across repetitions using the z-score of the Rand index (z-Rand). We summarized this measure by its mean and variance per γ. We then built an agreement matrix (pairwise co-assignment probability across repetitions) and derived a consensus partition using an iterative procedure^142^, with the consensus threshold set to the mean agreement from a label-shuffled null (node-wise permutations within runs). Optimal partitions were defined as those lying at stability peaks (high mean, low variance of z-Rand) within the refined sweep. When multiple local optima were present at distinct granularities, we retained all such solutions for downstream analyses and visualization. Note that singleton communities (<2 nodes) were removed for visualization only (rendered as NaN, remaining labels renumbered), and all quantitative similarity metrics were computed on the full label vectors.

#### Cross-partition similarity and contingency

Between-partition similarity was assessed with complementary, label-invariant measures: Adjusted Mutual Information (AMI) as a measure of primary similarity, and Variation of Information (VI) as a true distance metric. VI was used as both a distance for hierarchical clustering (average linkage) and dendrogram ordering of partitions, as well as inverted and scaled by its max (1 – VI / max(VI)) for use as a similarity measure to complement AMI. To interpret community structure relative to canonical systems, we compared each partition to the 7-network Yeo atlas. We constructed a contingency table (rows = Yeo systems, columns = partition communities) and computed both row-normalized (coverage of Yeo: “where did each Yeo system go?”; rows sum to 1) and column-normalized (purity of discovered communities with respect to Yeo; columns sum to 1) contingency. For visualization, we aligned columns to rows via a Hungarian (Kuhn-Munkres linear assignment) match that maximized overlap, however, all label-invariant metrics (AMI, VI) were computed before any matching. These analyses were performed using tools from the Brain Connectivity Toolbox (*community_louvain.m*, *agreement.m*, *consensus_und.m*) and a standard implementation of the Kuhn-Munkres assignment algorithm (*munkres.m*) in MATLAB.

### Communication modeling

We derived six graph-theoretic communication models from both weighted and binary structural connectomes to quantify alternative principles by which anatomical architecture may support large-scale neural interactions. These models were chosen to cover the routing-diffusion spectrum^19^ of communication processes: shortest-path efficiency(SPE), navigation efficiency (NE), search information efficiency (SIE), path transitivity (PT), communicability (CMY), and diffusion efficiency (DE). All models were computed from undirected connectivity matrices using established algorithms implemented in the Brain Connectivity Toolbox.

#### Weight vs cost network representations

Graph-theoretic communication models require both a weight representation of the connectome (reflecting the strength or capacity of an edge) and a cost or length representation (reflecting the difficulty or distance of traversing that edge). These representations are complementary: routing-based models (e.g., SPE) operate on costs, whereas diffusion-based models (e.g., CMY) operate on weights. A strong connection (high weight) should correspond to a short traversal cost, whereas weaker or absent connections should impose a high cost. In this study, each structural weighting was converted into an appropriate pair of matrices—one emphasizing capacity (weight) and the other emphasizing traversal difficulty (cost). Delay was treated directly as a cost (longer delays = larger costs), and its inverse (1/delay) served as the corresponding weight representation. Caliber- and myelin-weighted networks were treated as weights, and their cost counterparts were obtained using a monotonic transformation (L = −log W), ensuring that stronger connections map to shorter effective lengths. The log transform was chosen here in order to account for the highly skewed distribution of caliber edge weights. Binary connectivity was handled separately: the binary matrix itself served as the weight representation, whereas its cost version assigned a unit length to all existing edges and infinite length to absent edges, ensuring that shortest-path algorithms could only traverse anatomically present connections. This framework ensured internally consistent weight–cost pairs for each network variant and allowed all communication models to be computed in a uniform, interpretable manner.

#### Model implementations

SPE(Watts & Strogatz, 1998) was computed as the inverse of weighted shortest-path distance estimated using Floyd–Warshall searches. NE^143^ measured the success of greedy routing constrained by Euclidean embedding and was defined as the inverse of navigation path length. Search information^4,144–146^ quantified how deterministic shortest paths are from the perspective of a random walker. We used the inverse of search information as an efficiency-like measure, SIE. PT^4^ indexed the density of triangular detours along shortest paths. CMY(Crofts & Higham, 2009; Estrada & Hatano, 2008) was estimated using the matrix exponential of the degree-normalized adjacency matrix capturing the strength of multi-step walks between nodes. Let *A* be the weight matrix and *S* = diag(*A*1*) be the strength (weighted-degree) matrix. Then the normalized adjacency is *Ã* = *S*^−1⁄2^*AS*^−1⁄2^, and *CMY* = exp(*βÃ*). *β* was set to a value of 1 in this study to strike a balance between local and global communication patterns. DE^147–150^ was derived from mean first-passage times of a random walk and inverted to express greater diffusion efficiency as larger values.

All communication matrices were symmetrized, and the main diagonal was set to zero where appropriate. To support downstream regression modeling, each model additionally underwent a standardized preprocessing pipeline. For each matrix, skewness was evaluated; if the distribution exceeded a predefined skewness threshold (|skew| > 1.5), a log-transform was applied. All matrices were then z-scored across non-zero entries using a robust z-scaling function that preserves the distribution of meaningful values while ignoring structural zeros and infinities. Both raw and transformed versions of all communication models were retained for analysis, with the transformed variants used for all statistical modeling. This procedure yielded a consistent set of six communication models per structural network (SPE, NE, SIE, PT, CMY, DE), along with corresponding transformed versions.

### Spectral Analysis Framework

We developed a comprehensive spectral analysis framework to characterize scale-dependent network topology and to compare the structural organization of distinct microstructural weightings (caliber, MTsat, g-ratio, and delay). With this framework, we quantify the alignment, divergence, and scale distribution of topological features across networks.

#### Spectral decomposition and operators

All structural networks were first symmetrized and normalized to ensure comparability across weightings. Two complementary operators were analyzed:

1. the weighted adjacency matrix A, representing direct connection strength; and
2. the symmetrically normalized graph Laplacian L = I − D^−1/2^AD^−1/2^, representing diffusion- or flow-based topology. D is the diagonal strength matrix.

Each operator was decomposed into its full eigen-spectrum (eigenvalues λ_k_ and eigenvectors *v*_*k*_), which collectively describe the hierarchical modes of network organization—from global integrative patterns (low *k*) to fine-scale local motifs (high *k*).

#### Aggregate alignment with caliber

Before matching individual spectral components, we quantified the global alignment of each myelin-weighted network to the reference caliber network as the mean pairwise cosine similarity between corresponding eigenvectors:

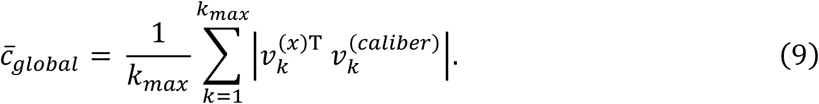

This provided an aggregate measure of spectral similarity that is insensitive to permutation of modes. To assess global shape differences in the eigenvalue distributions (spectral density), we also computed the Wasserstein (Earth Mover’s) distance between the normalized spectra of each network and the reference caliber network:

#### Windowed spectral subspaces

To facilitate scale-resolved comparisons, eigenvectors were partitioned into four contiguous k-windows that represent distinct topological scales:

1. **Global** (k = 2–20): large-scale integrative gradients.
2. **Meso** (k = 21–100): modular and subnetwork organization.
3. **Local** (k = 101–219): intermediate-scale motifs and dense clusters.
4. **Local**+ (k = 220–399): fine-grained, near-neighbor structure.

The boundaries of the four k-windows were determined algorithmically from the empirical distribution of caliber Laplacian eigenvector smoothness. Smoothness was quantified as the spatial variance of each eigenvector’s gradient across the cortical surface, then rank-ordered to form a cumulative distribution. The quantile inflection points of this distribution (approximately 0.05, 0.25, and 0.65) served as natural breakpoints, separating global, meso, local, and local+ scales. This procedure ensured that the window divisions reflected intrinsic transitions in the spatial frequency of caliber network topology rather than fixed k-intervals. Analyses were restricted to the non-trivial eigenmodes (k > 1), as the trivial first mode (k = 1) reflects only the connected structure of the graph. Each window was treated as a subspace representing a distinct topological regime.

#### Subspace alignment and similarity metrics

To assess structural correspondence across weightings, we implemented a windowed subspace alignment approach. For each k-window, the eigenvectors of each myelin-weighted network *A*_*x*_(or *L*_*x*_) were compared to those of the reference caliber-weighted network *A*_ref_ using cosine similarity between their corresponding subspaces. The mean cosine similarity quantified alignment strength, and Wasserstein distance between the eigenvalue distributions quantified spectral dissimilarity. These metrics were computed separately for adjacency and Laplacian operators, providing complementary views of topological correspondence based on raw connection strength (adjacency) and degree-normalized flow (Laplacian).

#### Topology of matched eigenvectors

To establish correspondence between eigenvectors of each target network and the caliber reference, we used a full-spectrum matching approach that incorporated a band-distance penalty to preserve spectral continuity. Cosine similarity was computed between every pair of eigenvectors across spectra, forming a similarity matrix. A penalty term proportional to the absolute difference in eigenvalue rank (|k_ref − k_tgt|) was subtracted from the similarity score, discouraging matches between eigenvectors that were widely separated in eigen-space. The resulting penalized cost matrix was then solved using the Hungarian (Munkres) algorithm, which identified the optimal one-to-one assignment minimizing total cost while maintaining local smoothness along the spectrum. This approach ensured robust matching across potential eigenvalue degeneracies and allowed continuity of correspondence across subspace boundaries without imposing hard window limits.

Within each k-window, matched eigenvectors were manually inspected, and a subset was selected for detailed topological analysis. Selection focused on those k-values where MTsat and g-ratio exhibited relatively poor alignment with caliber, whereas delay retained strong alignment. The intent was to apply region-of-interest logic to the spectral domain to isolate eigenmodes underpinning the divergence between myelin- and caliber-based topology. In an alternate stream, we averaged topology metrics across all eigenvectors within each window, which largely obscured our reported effects, confirming that the observed divergence arises from a restricted subset of spectral components.

On the subset of manually selected matched eigenvectors within each k-window, we computed Δ-metrics (x – ref) representing the difference between target and reference values for several topology descriptors:

- Modular fit (modR²): for each eigenvector *v*_*k*_, the proportion of variance in its node loadings explained by the Yeo-7 RSN assignments was computed via a one-way ANOVA, 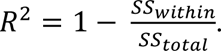 Higher modR² values indicate stronger modular alignment of the spectral mode with canonical functional systems.
- Participation ratio (PR): defined as 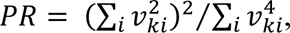 reflecting the effective number of nodes contributing to each eigenvector. Larger PR values correspond to more spatially distributed or cross-modular modes.
- Smoothness: quantified as the mean squared gradient of *v*_*k*_ across surface vertices, providing a continuous analog of spatial frequency.
- Moran’s I: spatial autocorrelation of *v*_*k*_across geodesic distance, used as a complementary measure to smoothness.
- S-A axis loading (SAr): Pearson correlation between *v*_*k*_and the cortical sensory-association axis, indicating the relative weighting of unimodal versus heteromodal systems.

All metrics were standardized (z-scored) within each network prior to computing Δ-values relative to caliber. All Δ-metrics were summarized (mean) within each k-window (global, meso, local, local+), providing a multi-scale profile of how myelin-weighted topology deviates from delay and caliber.

#### Spectral fingerprint analysis

To examine the overall distribution of topological “energy” across the eigen-spectrum, we computed spectral fingerprints for each communication model and weighting scheme. For each model, the communication matrix was projected onto the eigenbasis of its corresponding structural operator (adjacency or Laplacian). The squared projection coefficients, |*v*_*k*_^*T*^*Av*_*k*_ |^2^, were normalized by the total spectral energy to yield fractional energy distributions across eigenmodes. Fractions were then aggregated within each k-window to quantify the relative contribution of global, meso, local, and local+ topological scales to overall communication.

To ensure adequate representation across the spectrum, we evaluated coverage, defined as the proportion of total spectral energy captured by the analyzed eigenmodes. Because the first eigenvector (*k* = 1) accounted for a disproportionate share of the total energy—particularly for diffusion efficiency (≈0.99 for caliber; ≈0.95 for myelin) and for routing models under myelin weighting (≈0.8)—it was treated separately.

### Hierarchical Regression Framework

A hierarchical regression framework was implemented to quantify the unique contribution of myelin-weighted communication to the prediction of functional connectivity (FC). Analyses were performed at three spatial resolutions: global (whole-brain), pairwise resting-state network (RSN) interactions, and node-level. A nested model structure was used with ΔR² as the primary outcome metric and significance evaluated via nested F-tests (*α* < 0.05). Non-significant ΔR² values were excluded from subsequent analyses and visualizations. For each RSN pair, a sensitivity index was estimated by permuting model residuals (Freedman-Lane style^151^) to generate a null ΔR² distribution, against which true ΔR² values were z-scored (Fig. S8). The alignment of node-level ΔR² maps with the cortical sensory-association axis was quantified and significance was assessed using spin tests (n = 10,000, *α* < 0.05, hemisphere-constrained).

#### Individual myelin predictors

The first stage focused on quantifying how distinct myelin measures (MTsat, g-ratio and delay) contribute to the prediction of FC within individual communication models. SPE, NE, CMY, and DE models were tested. For each communication model (*CM_i_*), a hierarchical three-level design was applied: Level 1 included each myelin predictor individually (e.g., MTsat SPE); Level 2 included all pairwise combinations of the three myelin predictors; Level 3 included all three predictors simultaneously. Each level was tested relative to a fixed baseline model that included the structural backbone predictors: binary *CM_i_*, caliber *CM_i_*, and Euclidean distance. Thus, all ΔR² values reflect improvement in explained variance relative to this base model rather than incremental changes across levels.

#### Cross-Model and Interaction Effects

The second stage shifted focus from individual myelin measures to the communication models themselves, i.e., all models considered all myelin metrics combined. The design systematically examined how routing-type and diffusion-type communication models—both separately and together—contribute to FC prediction. We further tested whether these effects were modified by interactions with myelin communication. Five distinct interaction blocks were tested: (1) main effects only (no interactions); (2) interactions between caliber and myelin communication; (3) interactions between myelin communication and Euclidean distance; (4) the combined interactions of myelin communication with both caliber communication and Euclidean distance; and (5) the total contribution of myelin communication (main effects + interactions). For each block, two levels were evaluated: level 1 tested routing models (e.g., SPE, NE) and diffusion models (e.g., DE, CMY) separately; and level 2 tested routing and diffusion models jointly. Each model comparison was performed relative to its own base model, which always included Euclidean distance, caliber, and binary connectivity for the relevant communication model or model pair. ΔR² again represented the gain in explained variance relative to this base, however ΔR² of the interaction blocks (2-4) represented the *incremental gain* from adding interactions i.e., it was measured relative to the base model + the main effects.

#### Myelin vs caliber communication (ΔΔ*R*^2^)

To directly compare the predictive contributions of myelin- versus caliber-weighted communication, we computed a differential effect size, ΔΔR², defined as the difference in ΔR² obtained when replacing the caliber-weighted communication model with its myelin-weighted counterpart while holding all other predictors constant. For a given communication model (e.g., SPE), two matched hierarchical regressions were fit: one using the caliber-weighted communication matrix and one using the corresponding myelin-weighted matrix, each tested relative to an identical base model containing the binary communication matrix and Euclidean distance. The resulting ΔR² values quantify the unique gain in explained variance attributable to either caliber or myelin communication. Their difference, 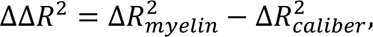 indexes whether myelin communication provides stronger, weaker, or equivalent explanatory power relative to caliber when entered into an otherwise comparable model. Positive values indicate a larger contribution from myelin-weighted communication.

#### Statistical implementation

Regression models were implemented using standard multiple linear regression with ordinary least squares estimation. All predictors were z-scored prior to entry into the model to ensure comparability of coefficients and stability of nested F-tests. Model residuals were verified to approximate normality, and variance-inflation diagnostics were used to confirm the effect of multicollinearity did not exceed a tolerance value of 10. When computing ΔR², F-statistics were derived from nested model comparisons following the standard formulation for non-orthogonal regressors. An elasticity analysis was also performed for each RSN-pair to probe the stability of model fit under small heterogeneous perturbations of predictor weights. Elasticity was computed by applying a row-wise dilation of 1%, 2.5%, and 5%, corresponding to proportional changes relative to each variable’s standard deviation. For node-wise analyses, regression and permutation tests were vectorized for computational efficiency, and spatial permutation (spin) tests used hemisphere-preserving rotations based on the fs_LR 32k surface geometry. The aggregation of summary statistics (ΔR², sensitivity index, elasticity) was performed within FC frequency bands before computing cross-band summaries.

## Acknowledgments

The authors acknowledge research support from the Natural Sciences and Engineering Research Council of Canada (NSERC), the Canadian Institutes for Health Research (CIHR), Fonds de recherche du Québec – Santé (FRQ-S), CFREF Healthy Brains for Healthy Lives, the Killam Trusts, and Brain Canada.

## Notes

### Competing Interest Statement

The authors have declared no competing interest.

