## Supplementary Information for "White Matter Myelin Shapes Macroscale Functional Connectivity Through Integrative Communication"

### Supplementary Material

Mark C. Nelson[1,2], Wen Da Lu[2,3], Ilana R. Leppert[2], Golia Shafiei[4,5,6], Heather A. Hansen[7], Christopher D. Rowley[8], Bratislav Misic[1,2], and Christine L. Tardif[1,2,3]

*[1]Department of Neurology and Neurosurgery, McGill university, Montreal, QC, Canada. [2]McConnell Brain Imaging Centre, Montreal Neurological Institute and Hospital, Montreal, QC, Canada. [3]Department of Biomedical Engineering, McGill University, Montreal, QC, Canada. [4]Penn Lifespan Informatics and Neuroimaging Center (PennLINC), Perelman School of Medicine, University of Pennsylvania, Philadelphia, PA, USA. [5]Department of Psychiatry, Perelman School of Medicine, University of Pennsylvania, Philadelphia, PA, USA. [6]Lifespan Brain Institute (LiBI), Children's Hospital of Philadelphia and Penn Medicine, Philadelphia, PA, USA. [7]Department of Psychological Sciences, William & Mary University, Williamsburg, VA, USA. [8]Department of Physics and Astronomy, McMaster University, Hamilton, ON, Canada.*

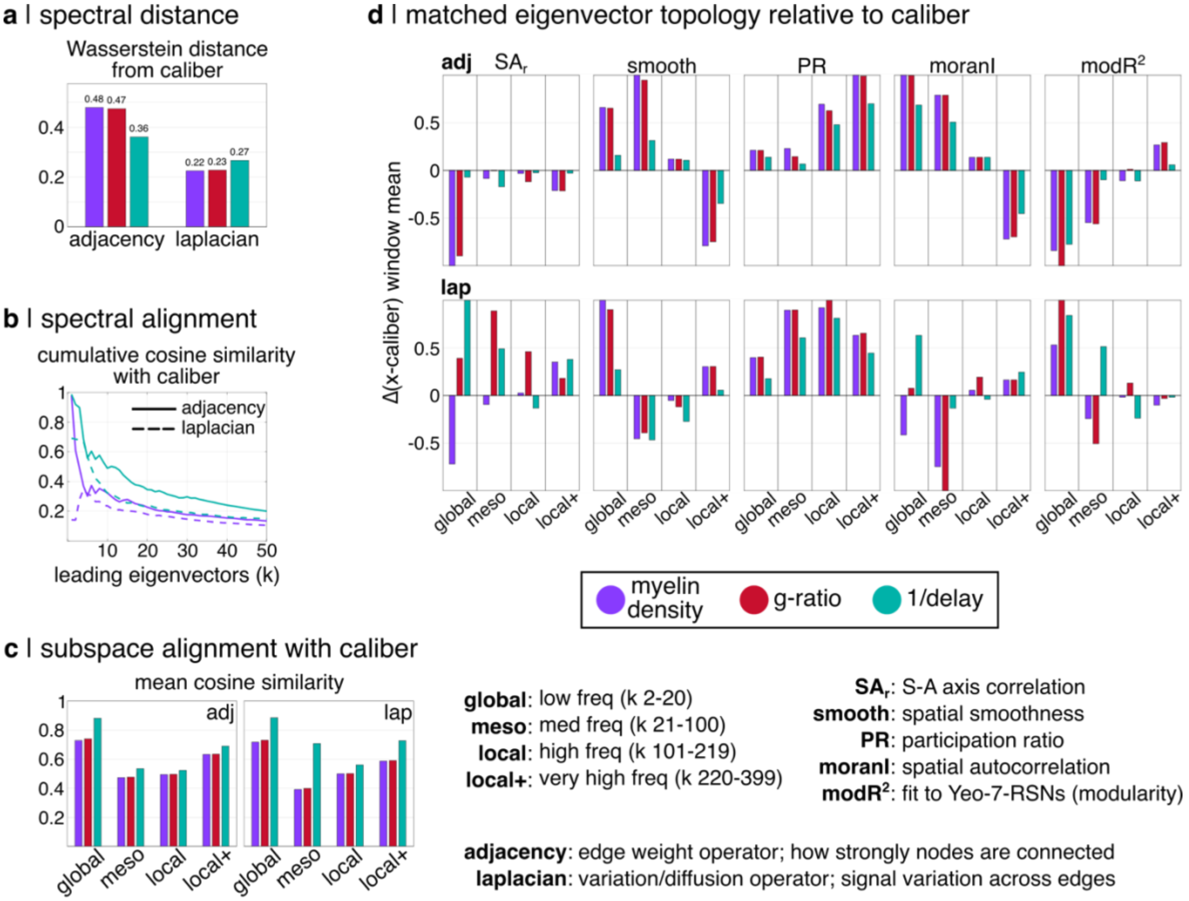


*Figure S1.* ***Spectral comparison of myelin-sensitive connectivity to tract caliber connectivity***. *(a) The spectral distance from caliber is quantified as the Wasserstein distance (earth mover's distance) between eigenvalue distributions. (b) The spectral alignment with caliber is quantified as the cumulative mean cosine similarity between eigenvectors (running mean from 1 to k). (c) The spectral alignment with caliber is shown within windowed subspaces corresponding to distinct topological scales. Eigenvectors were matched between target and reference networks using Hungarian matching, and mean cosine similarity is reported for each window. (d) Network topology is compared to caliber across a manually selected subset of matched eigenvectors from each windowed subspace. Eigenvectors of interest for each subspace are chosen to accentuate relative alignment of 1/delay and relative misalignment of myelin density (MTsat) and g-ratio. Reported values reflect the window mean of Δ values (myelin – caliber). Both the adjacency and Laplacian operators are examined in all panels. Only the first 50 eigenvectors are considered for panels (a) and (b), whereas the full eigenspectrum is considered in panels (c) and (d). The 1$^{st}$ trivial eigenmode (k = 1) is excluded for the Laplacian operator in all cases.*

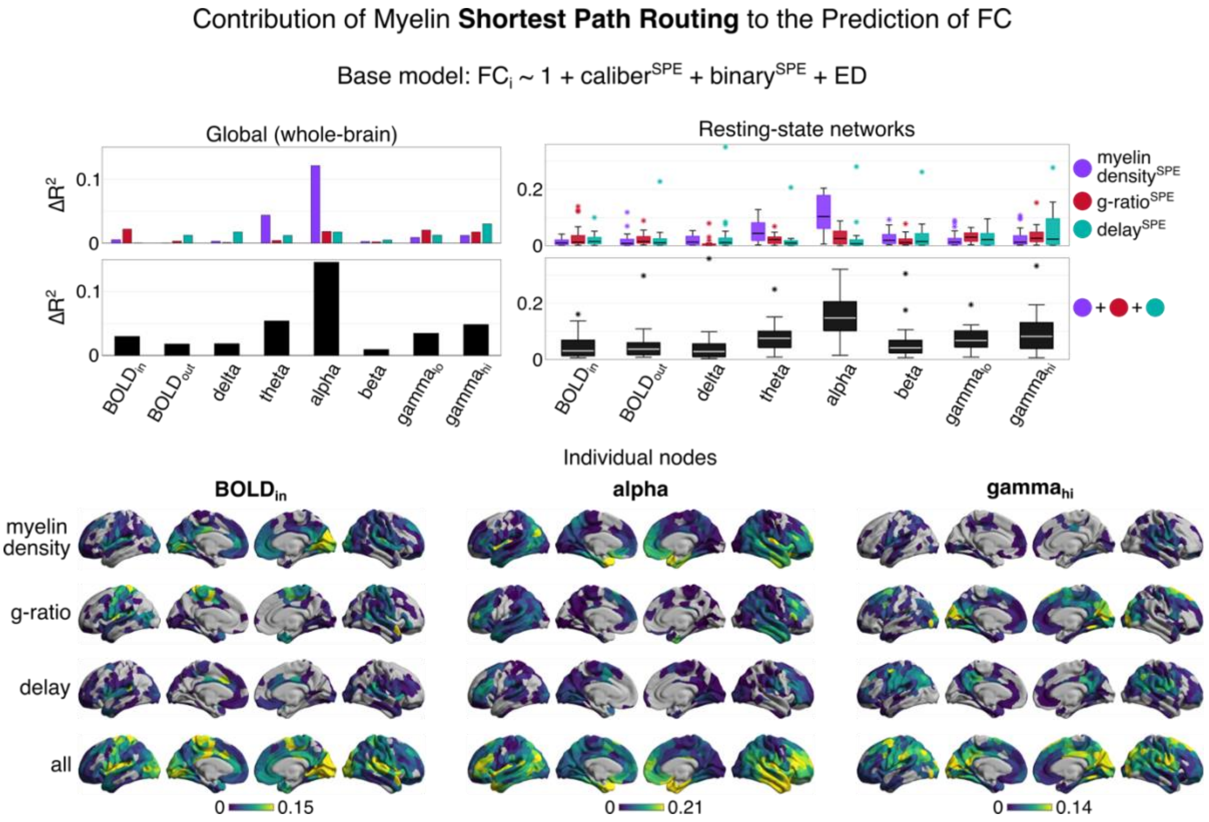


*Figure S2.* ***The contribution of myelin-sensitive shortest path routing to the prediction of functional connectivity (FC)****. The coupling of myelin-sensitive communication and FC is assessed using nested regression at three spatial scales: global (whole-brain), pairwise resting-state networks, and individual nodes. Two conditions are tested: (i) individual myelin-sensitive predictors, and (ii) all myelin-sensitive predictors combined. The reported* $\Delta R^2$ *values reflect the contribution of myelin-sensitive communication relative to the base model. The base regression model contains communication measures computed from edge caliber and binary networks, as well as the Euclidean distance (ED) between node pairs. Nested F-tests are used to filter non-significant values (α = 0.05). The Schaefer-400 parcellation defines cortical nodes and the Yeo-7-Network atlas defines resting-state networks. Boxplots contain standard components. The shortest path efficiency (SPE) communication model is shown.*

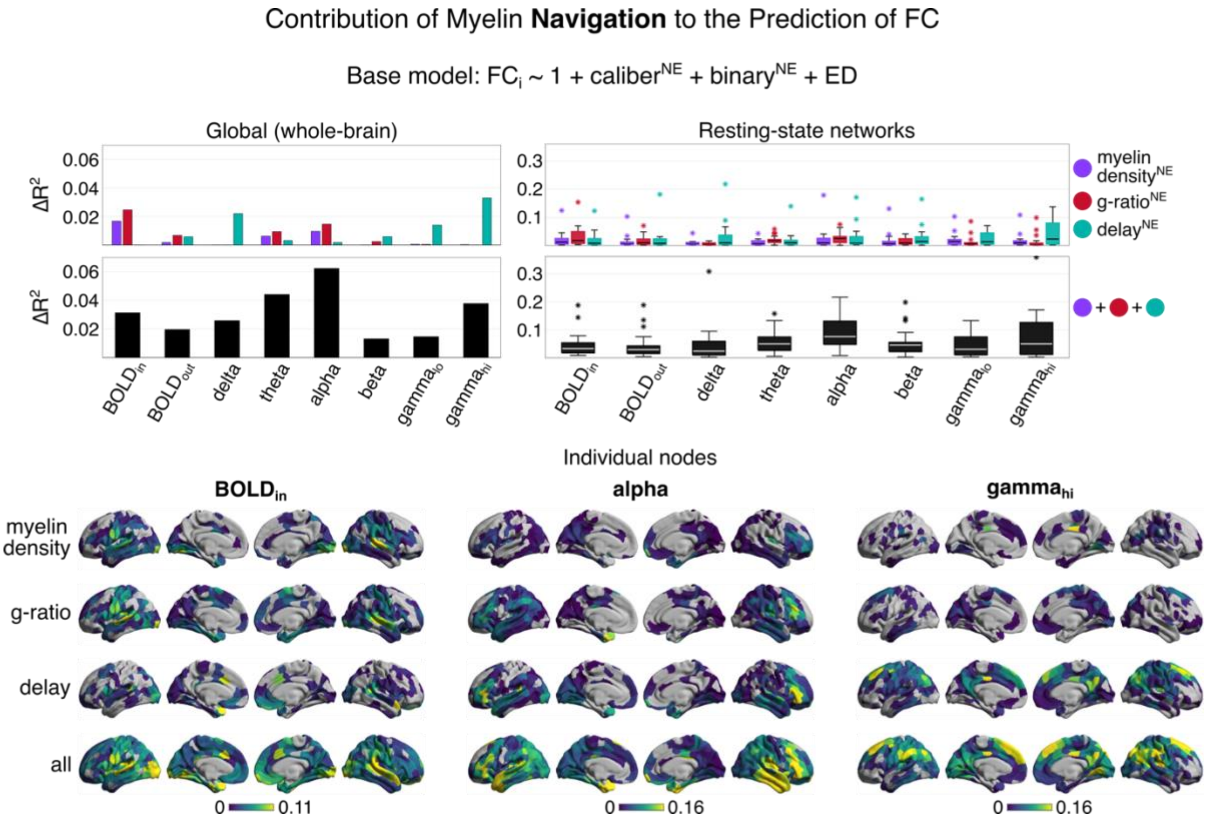


*Figure S3*. ***The contribution of myelin-sensitive navigation to the prediction of functional connectivity (FC)***. *The coupling of myelin-sensitive communication and FC is assessed using nested regression at three spatial scales: global (whole-brain), pairwise resting-state networks, and individual nodes. Two conditions are tested: (i) individual myelin-sensitive predictors, and (ii) all myelin-sensitive predictors combined. The reported* $\Delta R^2$ *values reflect the contribution of myelin-sensitive communication relative to the base model. The base regression model contains communication measures computed from edge caliber and binary networks, as well as the Euclidean distance (ED) between node pairs. Nested F-tests are used to filter non-significant values (*$\alpha = 0.05$*). The Schaefer-400 parcellation defines cortical nodes and the Yeo-7-Network atlas defines resting-state networks. Boxplots contain standard components. The navigation efficiency (NE) communication model is shown.*

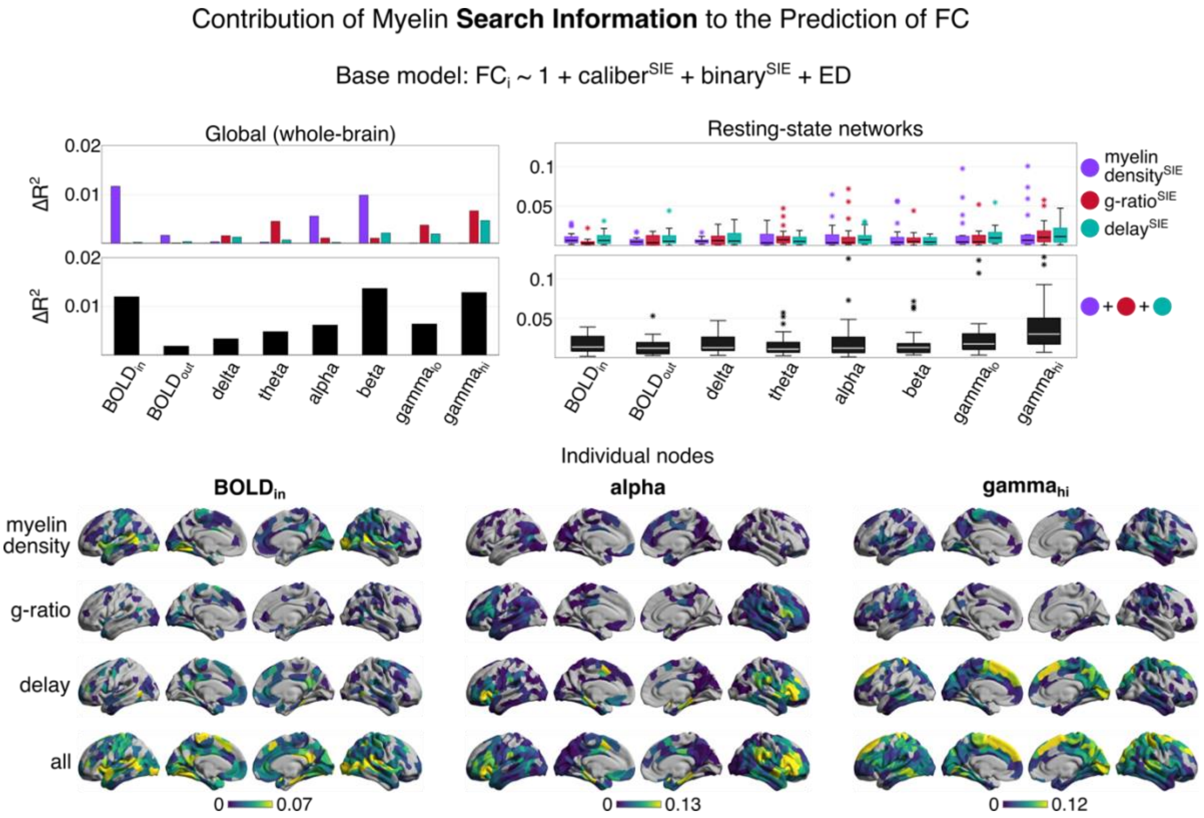


*Figure S4.* ***The contribution of myelin-sensitive search information to the prediction of functional connectivity (FC)***. *The coupling of myelin-sensitive communication and FC is assessed using nested regression at three spatial scales: global (whole-brain), pairwise resting-state networks, and individual nodes. Two conditions are tested: (i) individual myelin-sensitive predictors, and (ii) all myelin-sensitive predictors combined. The reported $\Delta R^2$ values reflect the contribution of myelin-sensitive communication relative to the base model. The base regression model contains communication measures computed from edge caliber and binary networks, as well as the Euclidean distance (ED) between node pairs. Nested F-tests are used to filter non-significant values ($\alpha = 0.05$). The Schaefer-400 parcellation defines cortical nodes and the Yeo-7-Network atlas defines resting-state networks. Boxplots contain standard components. The search information efficiency (SIE) communication model is shown.*

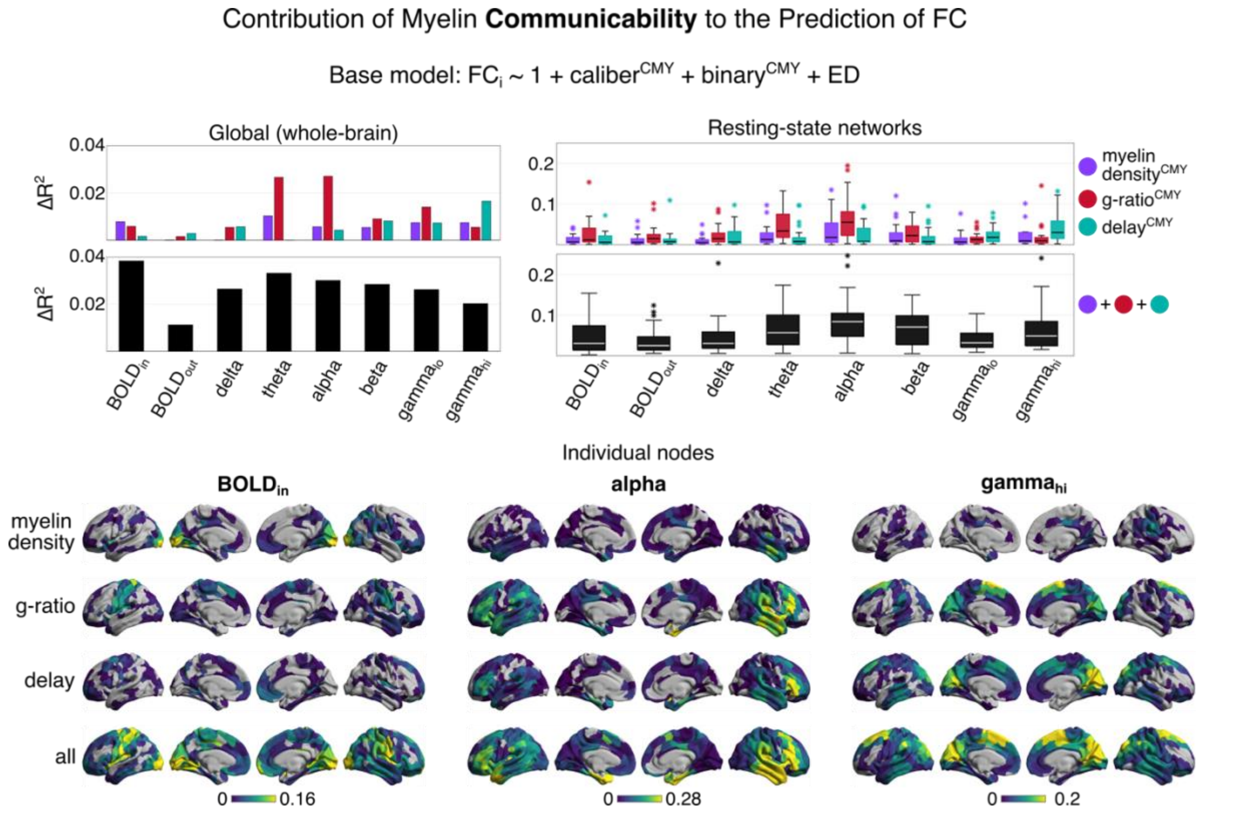


*Figure S5.* ***The contribution of myelin-sensitive communicability to the prediction of functional connectivity (FC)****. The coupling of myelin-sensitive communication and FC is assessed using nested regression at three spatial scales: global (whole-brain), pairwise resting-state networks, and individual nodes. Two conditions are tested: (i) individual myelin-sensitive predictors, and (ii) all myelin-sensitive predictors combined. The reported $\Delta R^2$ values reflect the contribution of myelin-sensitive communication relative to the base model. The base regression model contains communication measures computed from edge caliber and binary networks, as well as the Euclidean distance (ED) between node pairs. Nested F-tests are used to filter non-significant values ($\alpha = 0.05$). The Schaefer-400 parcellation defines cortical nodes and the Yeo-7-Network atlas defines resting-state networks. Boxplots contain standard components. The communicability (CMY) communication model is shown.*

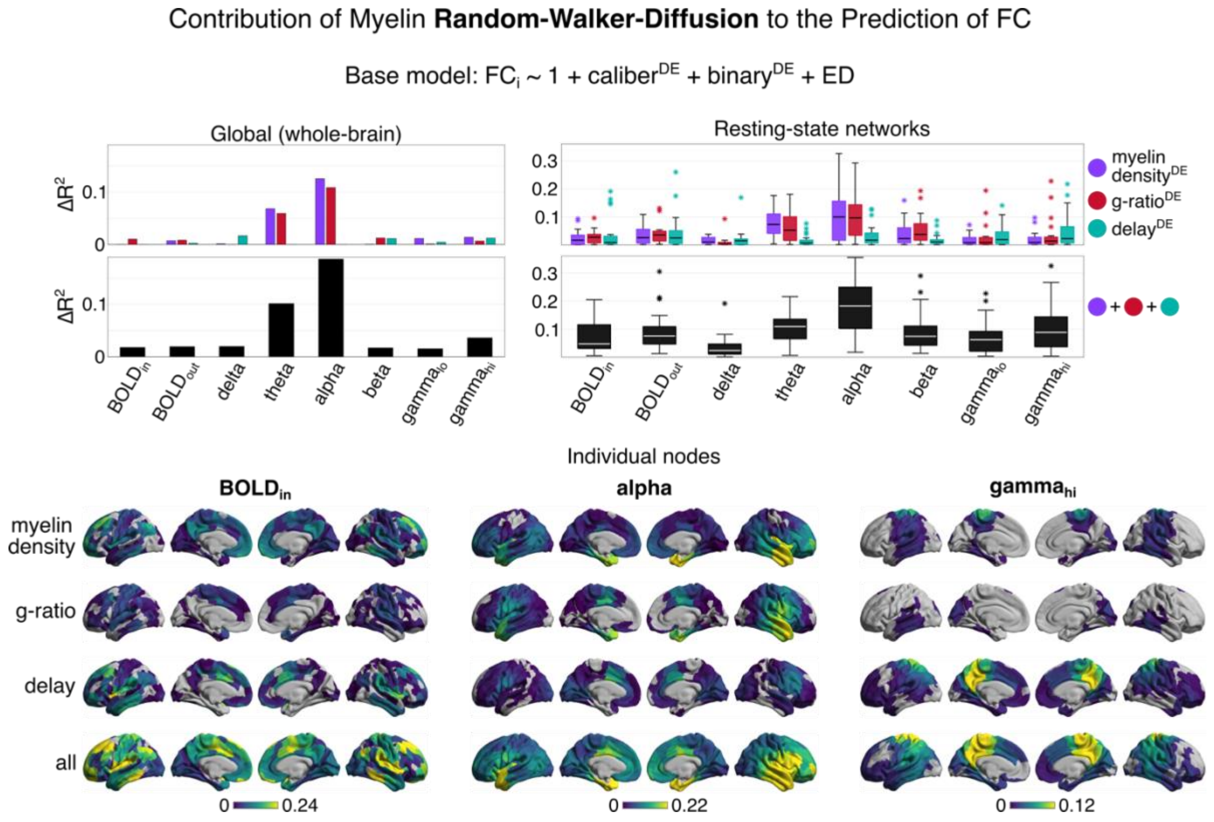


*Figure S6.* ***The contribution of myelin-sensitive random-walker diffusion to the prediction of functional connectivity (FC)****. The coupling of myelin-sensitive communication and FC is assessed using nested regression at three spatial scales: global (whole-brain), pairwise resting-state networks, and individual nodes. Two conditions are tested: (i) individual myelin-sensitive predictors, and (ii) all myelin-sensitive predictors combined. The reported* $\Delta R^2$ *values reflect the contribution of myelin-sensitive communication relative to the base model. The base regression model contains communication measures computed from edge caliber and binary networks, as well as the Euclidean distance (ED) between node pairs. Nested F-tests are used to filter non-significant values (α = 0.05). The Schaefer-400 parcellation defines cortical nodes and the Yeo-7-Network atlas defines resting-state networks. Boxplots contain standard components. The diffusion efficiency (DE) communication model is shown.*

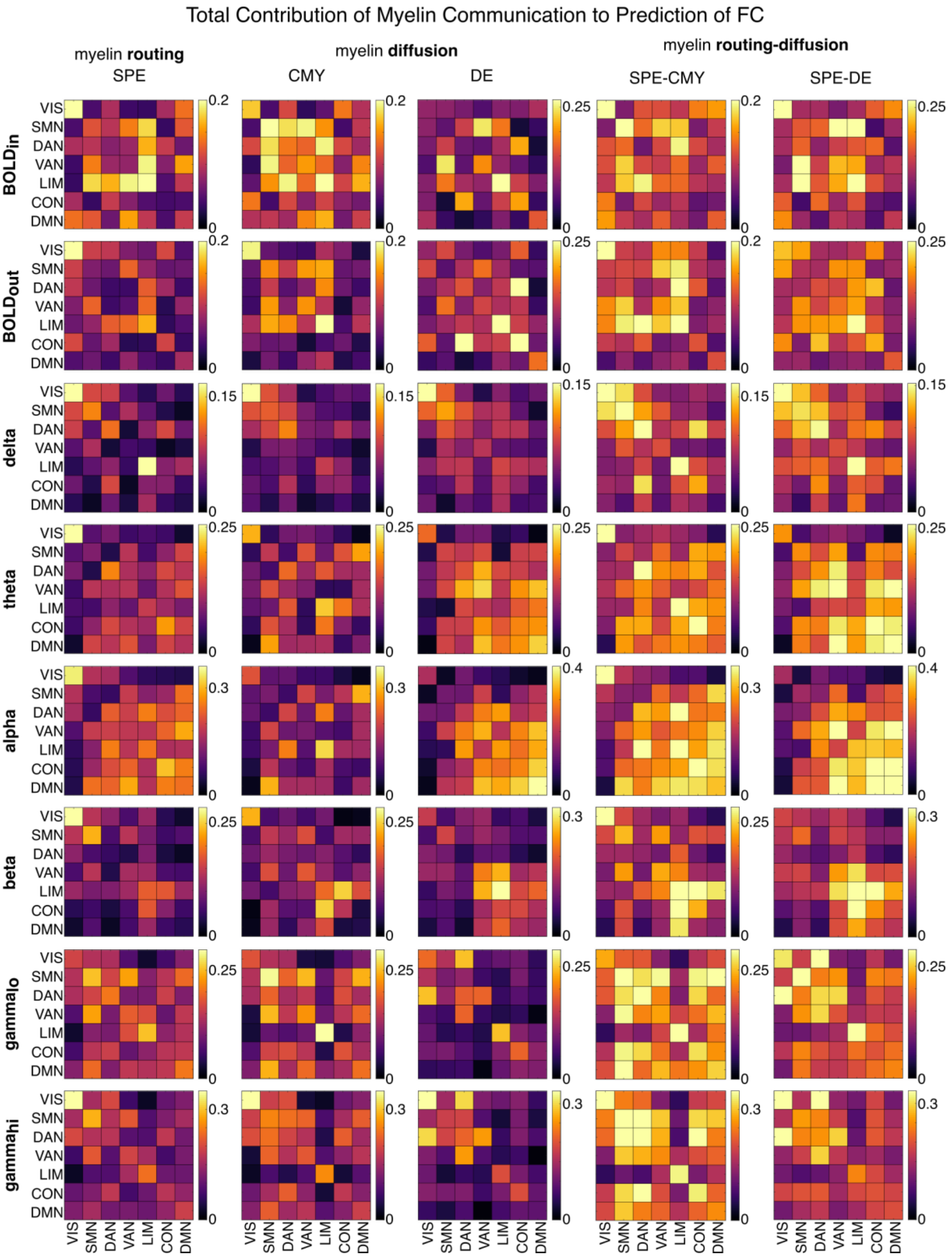


*Figure S7*. ***The total contribution of myelin-sensitive communication to the prediction of functional connectivity (FC)***. *The coupling of myelin-sensitive communication and FC is assessed using nested*

*regression across pairs of resting-state networks. All models feature all three myelin-sensitive predictors (myelin density, g-ratio, and delay). The reported* $\Delta R^2$ *values reflect the contribution of myelin-sensitive communication relative to the base model. The base regression model contains communication measures computed from edge caliber and binary networks, as well as the Euclidean distance (ED) between node pairs. Nested F-tests are used to filter non-significant values (*$\alpha = 0.05$*). The Schaefer-400 parcellation defines cortical nodes and the Yeo-7-Network atlas defines resting-state networks. Resting-state networks correspond to: visual (VIS), somatomotor (SMN), dorsal attention (DAN), ventral attention (VAN), limbic (LIM), fronto-parietal control (CON), and default mode (DMN). Communication models correspond to: shortest path efficiency (SPE), communicability (CMY), and diffusion efficiency (DE).*

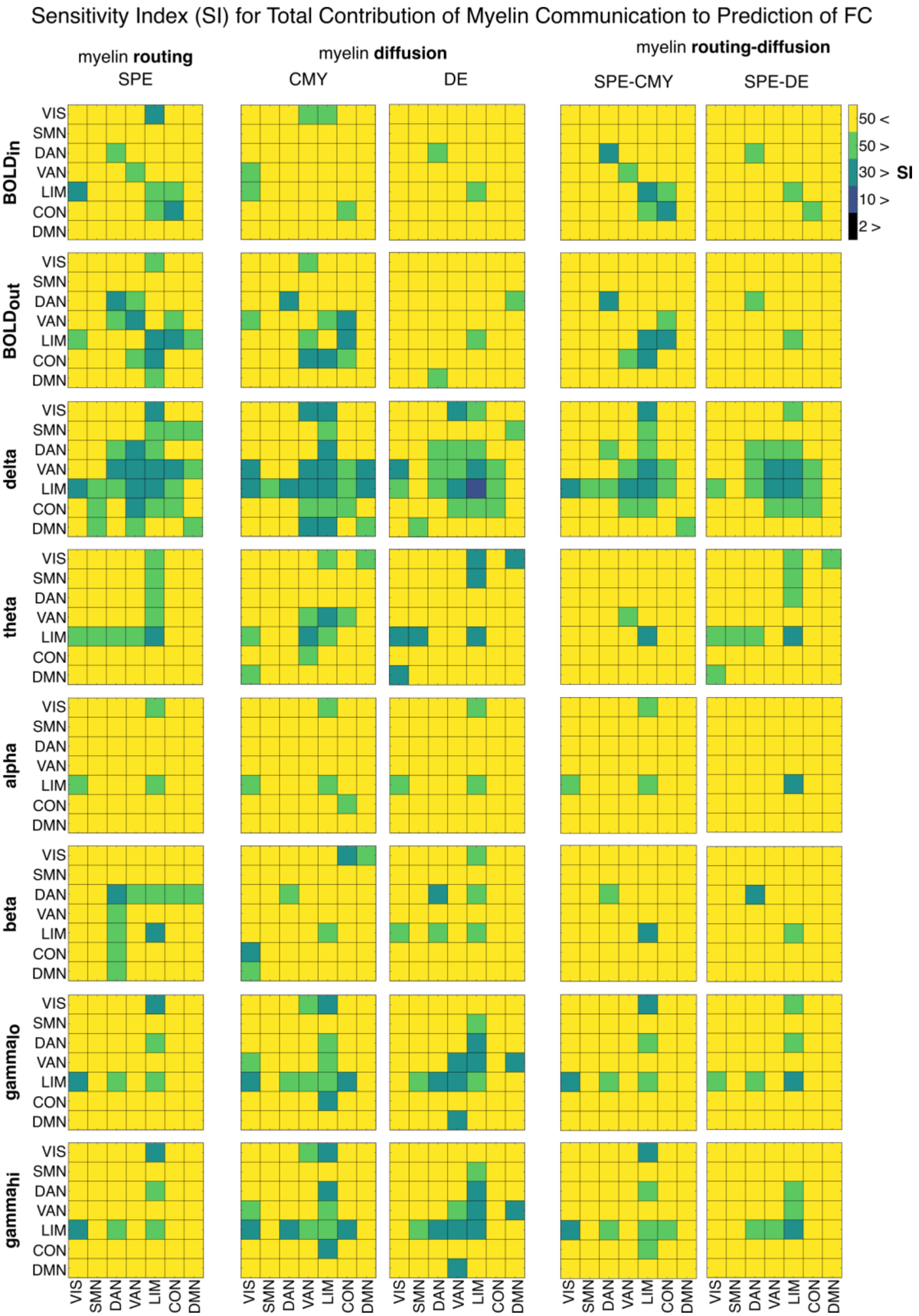


*Figure S8.* ***The significance of coupling between myelin-sensitive communication and functional connectivity (FC)****. A sensitivity index (SI) value is shown for each resting-state network pair. SI is*

*estimated by permuting nested-regression model residuals to obtain a null $\Delta R^2$ distribution against which the true $\Delta R^2$ values are z-scored. All models feature all three myelin-sensitive predictors (myelin density, g-ratio, and delay). The true $\Delta R^2$ values reflect the contribution of myelin-sensitive communication relative to the base model. The base regression model contains communication measures computed from edge caliber and binary networks, as well as the Euclidean distance (ED) between node pairs. Nested F-tests are used to filter non-significant values ($\alpha = 0.05$). The Schaefer-400 parcellation defines cortical nodes and the Yeo-7-Network atlas defines resting-state networks. Resting-state networks correspond to: visual (VIS), somatomotor (SMN), dorsal attention (DAN), ventral attention (VAN), limbic (LIM), fronto-parietal control (CON), and default mode (DMN). Communication models correspond to: shortest path efficiency (SPE), communicability (CMY), and diffusion efficiency (DE).*

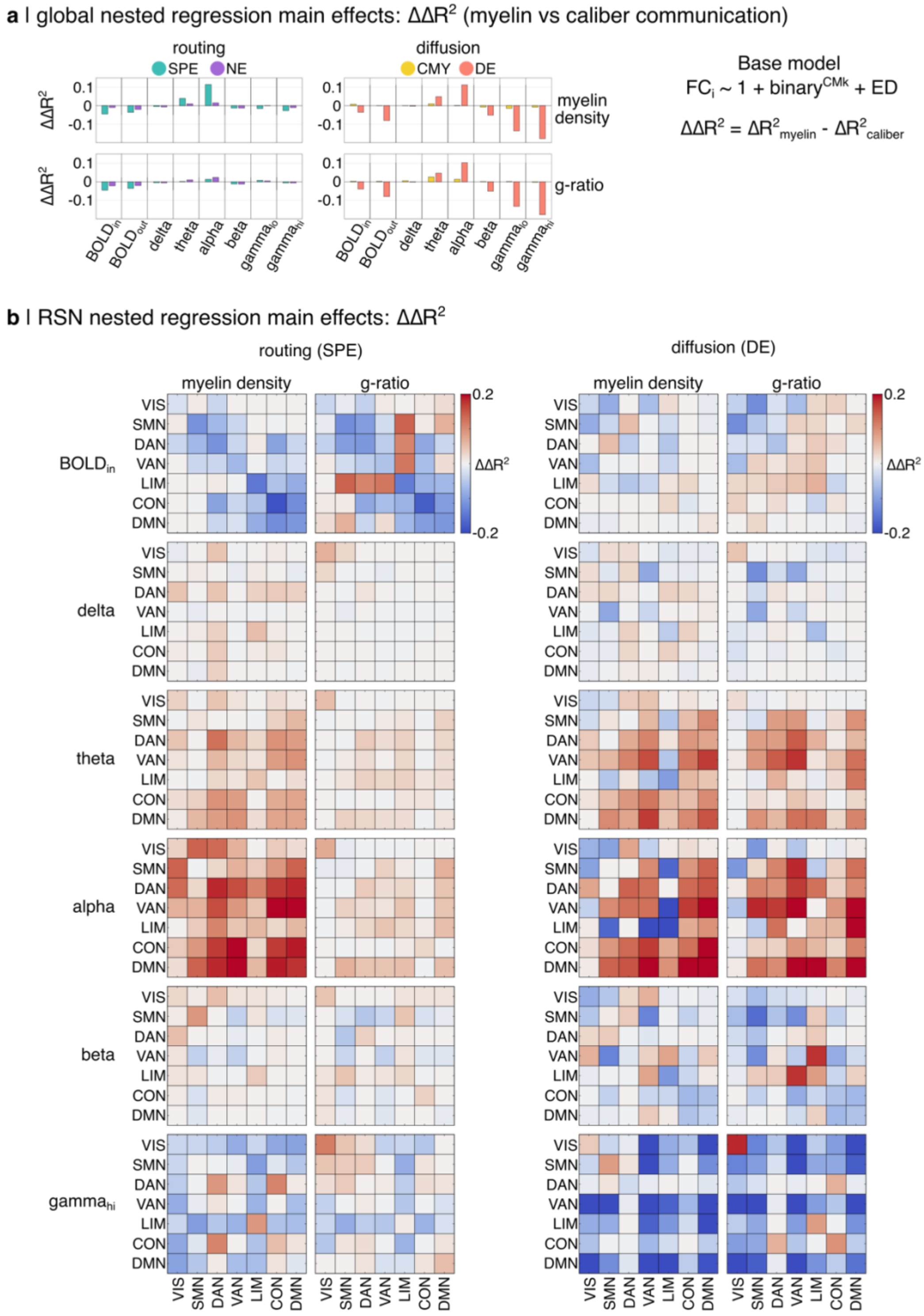


*Figure S9.* ***The relative magnitude of functional coupling for myelin-sensitive vs caliber-sensitive communication****. The strength of functional coupling is compared between myelin-sensitive and caliber-sensitive communication using nested regression at the (a) global and (b) pairwise resting-state network levels. The reported $\Delta\Delta R^2$ values reflect the difference between separate nested regression models testing*

*the contribution of myelin-sensitive and caliber-sensitive communication ($\Delta\Delta R^2 = \Delta R^2_{myelin} - \Delta R^2_{caliber}$). Positive $\Delta\Delta R^2$ values indicate stronger functional coupling for myelin-sensitive communication. All $\Delta R^2_{myelin}$ and $\Delta R^2_{caliber}$ values reflect the gain in model fit relative to the same base model. The base regression model contains communication measures computed from a binary network and the Euclidean distance (ED) between node pairs. All myelin-sensitive communication measures are derived using a single myelin-sensitive network (myelin density or g-ratio). Edge caliber is derived using the COMMIT framework and quantifies the total intra-axonal cross-sectional area of the streamlines comprising each network edge. Nested F-tests are used to filter non-significant values ($\alpha = 0.05$). The Schaefer-400 parcellation defines cortical nodes and the Yeo-7-Network atlas defines resting-state networks. Resting-state networks correspond to: visual (VIS), somatomotor (SMN), dorsal attention (DAN), ventral attention (VAN), limbic (LIM), fronto-parietal control (CON), and default mode (DMN). Communication models correspond to: shortest path efficiency (SPE), navigation efficiency (NE), communicability (CMY), and diffusion efficiency (DE).*

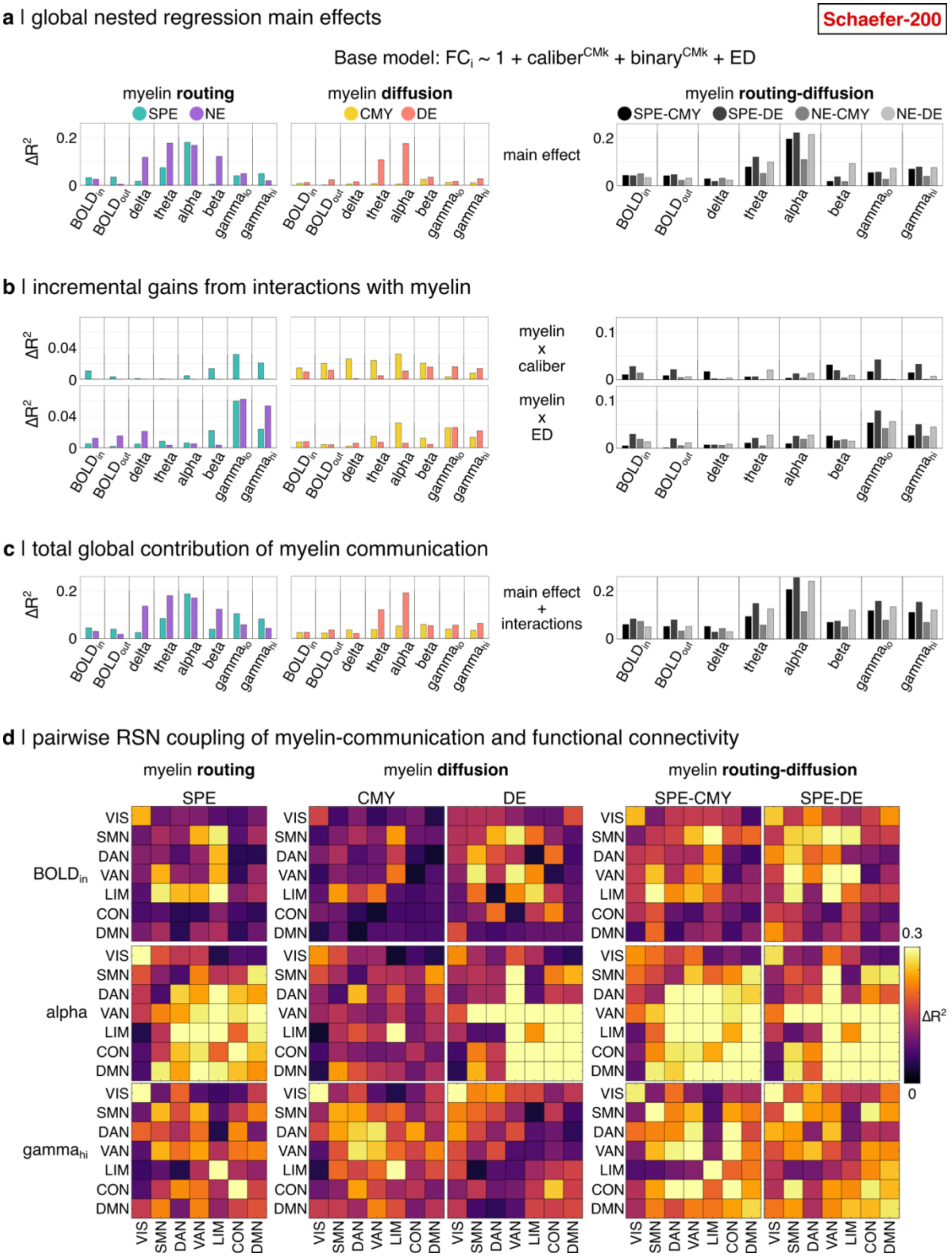


*Figure S10. **Replication of main regression results using Schaefer-200**. The coupling of myelin-sensitive communication and functional connectivity is assessed using nested regression at two spatial scales. At the global level, (a) the main effects, (b) the incremental gains from interactions, and (c) the total*

*contribution (main effects + interactions) of myelin communication are shown. (d) The total contribution of myelin communication is shown for all pairs of resting-state networks. All models feature all three myelin-sensitive predictors (myelin density, g-ratio, and delay). The reported $\Delta R^2$ values reflect the contribution of myelin-sensitive communication relative to the base model. The base regression model contains communication measures computed from edge caliber and binary networks, as well as the Euclidean distance (ED) between node pairs. Nested F-tests are used to filter non-significant values ($\alpha$ = 0.05). The Schaefer-200 parcellation defines cortical nodes and the Yeo-7-Network atlas defines resting-state networks. Resting-state networks correspond to: visual (VIS), somatomotor (SMN), dorsal attention (DAN), ventral attention (VAN), limbic (LIM), fronto-parietal control (CON), and default mode (DMN). Communication models correspond to: shortest path efficiency (SPE), navigation efficiency (NE), communicability (CMY), and diffusion efficiency (DE).*

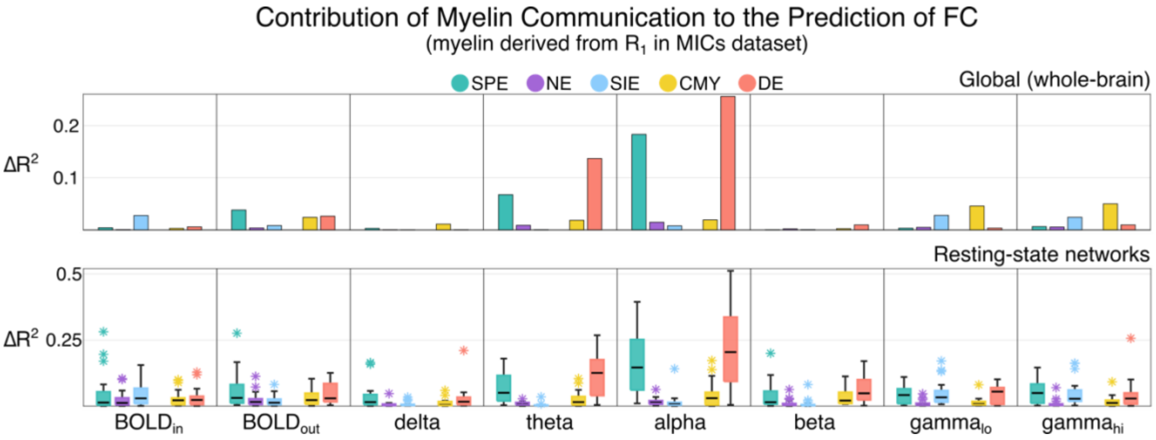


*Figure S11.* ***Testing coupling of myelin-sensitive communication and functional connectivity (FC) using independent dataset****. The coupling of myelin-sensitive communication and FC is assessed using nested regression at the global and pairwise resting-state network levels. Structural and functional data are derived from an independent multi-modal dataset ([https://osf.io/j532r/overview](https://osf.io/j532r/overview)). The myelin-sensitive metric in these data corresponds to the $R_1$ relaxation rate ($1/T_1$). The reported $\Delta R^2$ values reflect the contribution of myelin-sensitive communication relative to the base model. The base regression model contains communication measures computed from edge caliber and binary networks, as well as the Euclidean distance (ED) between node pairs. Nested F-tests are used to filter non-significant values ($\alpha$ = 0.05). The Schaefer-400 parcellation defines cortical nodes and the Yeo-7-Network atlas defines resting-state networks. Boxplots contain standard components. Communication models correspond to: shortest path efficiency (SPE), navigation efficiency (NE), search information efficiency (SIE), communicability (CMY), and diffusion efficiency (DE).*
